# A longitudinal atlas of post-viral lung regeneration reveals persistent injury-associated cell states

**DOI:** 10.1101/2024.05.24.595801

**Authors:** Terren K. Niethamer, Joseph D. Planer, Michael P. Morley, Apoorva Babu, Maria C. Basil, Edward Cantu, David B. Frank, Joshua M. Diamond, Ana N. Nottingham, Shanru Li, Lillian I. Levin, Su Zhou, Edward E. Morrisey

**Author notes:** Address correspondence to: Edward E. Morrisey, Ph.D., University of Pennsylvania, Translational Research Center, Room 11-124, 3400 Civic Center Boulevard, Building 421, Philadelphia, PA 19104-5129; Terren K. Niethamer, Ph.D., Head, Regenerative Plasticity and Signaling Section, Cancer & Developmental Biology Laboratory (CDBL), Center for Cancer Research (CCR) – Frederick, National Cancer Institute (NCI), National Institutes of Health (NIH), 1050 Boyles Street, Building 560, Frederick, MD 21702. These authors contributed equally to this work (Terren K. Niethamer, Joseph D. Planer).

## Abstract

Functional regeneration of the lung’s gas exchange surface following injury requires the coordination of a complex series of cell behaviors within the alveolar niche. Using a multi-modal approach, we have mapped the temporal sequencing of mouse lung regeneration after acute viral injury, demonstrating that this response is asynchronously phased across different cellular compartments. This longitudinal atlas of regeneration has produced a catalogue of new cell states that reflect transient and persistent transcriptional alterations in daughter cells as they transit across axes of differentiation. These new cell states include an injury-induced capillary endothelial cell (iCAP) that arises after injury, persists indefinitely, and shares transcriptional hallmarks with both developing lung endothelium and the endothelial aberrations found in degenerative human lung diseases. This comprehensive atlas of lung regeneration provides a foundational resource to understand the complexity of the cellular and molecular responses to injury, reveals the critical importance of capillary endothelium in maintaining and rebuilding the alveolar niche after injury, and correlates these responses to those found in development and human lung diseases.

## Introduction

One of the major functions of the lung is gas exchange with the external environment, which occurs through the interface between the pulmonary epithelium lining alveoli and the capillary endothelium lining blood vessels in the distal lung. The epithelium and endothelium that comprise this interface share a fused basement membrane^1^, rendering them highly susceptible to injury. If the lung is damaged, repairing the alveolar niche to reestablish gas exchange is critical for restoring lung function. While regeneration of the alveolus may have characteristics in common with formation of the alveolar niche during development, there have been few studies to directly compare these processes. Mouse and human embryonic and postnatal lung development have been profiled^2–6^, and lung homeostasis and regeneration have also been studied in both mouse and human^7–11^. However, a longitudinal and long-term analysis of the response to lung injury across cellular compartments is lacking.

Unlike other organs with continuously regenerating epithelia, the lung is quiescent at homeostasis. After injury, facultative progenitor cells can be activated to repair the epithelium^12–15^. Proliferation of alveolar type 2 (AT2) cells and differentiation of AT2 cells into alveolar type 1 (AT1) cells are temporally and spatially regulated processes^16,17^. Preliminary evidence suggests that proliferation of pulmonary endothelial cells (ECs) does not necessarily parallel the temporal or spatial characteristics of epithelial proliferation. In mice with acute lung injury due to influenza infection, EC proliferation continues to increase into the third week following infection and is similar in magnitude across damaged zones and regions with seemingly normal tissue structure^9,18^. In addition, endothelial regeneration is necessary to re-form functional alveoli after injury, and factors affecting EC proliferation are essential in this process^19,20^.

Transitional cell states have been increasingly recognized after acute lung injury. AT2 cells adopt a damage-associated transient progenitor (DATP) or pre-alveolar type 1 transitional (PATS) cell state marked by *Krt8*/*Cldn4* expression in response to lung injury^21–23^. This transitional cell state is also characterized by increased expression of genes associated with DNA damage and senescence^22,23^, and could be induced by the intense inflammation that occurs during infectious lung injury^21^. Inflammatory signaling within the alveolus is a double-edged sword promoting either the restoration of homeostasis after lung injury^24,25^ or the persistence of damage-associated states that impede functional regeneration^21,26,27^. Whether non-epithelial cell types in the lung undergo similar transitions in cell state remains less clear.

Here, we use single cell transcriptomics combined with lineage tracing of proliferating progenitors to comprehensively define the long-term, longitudinal response of the mouse lung to influenza-induced acute injury and regeneration. Our results demonstrate that proliferative dynamics and cellular composition vary widely both across cellular compartments in the lung and within compartments across time, altering cell-cell communication as the lung regenerates. This lung regeneration atlas comprehensively catalogues intermediate or transitional cell states that arise after viral injury. Although some of these cellular states resolve, others persist indefinitely, suggesting that the lung does not readily return to its pre-injury condition. An injury-induced capillary (iCAP) state is identified in the pulmonary endothelium. iCAPs are preferentially found in regions of densely injured tissue after injury and arise independently of aberrant Krt5^+^ epithelium and interferon-γ (IFN-γ) signaling. Lineage tracing using novel CAP1- and CAP2-specific mouse lines shows that iCAPs are derived from both CAP1 and CAP2 ECs during regeneration. The iCAP state is characterized by high expression of *Sparcl1* and *Ntrk2,* and shares transcriptional similarities with ECs of other organs as well as with immature lung ECs present prior to the transition to air breathing. iCAP cells also possess an inflammatory transcriptional signature, which is observed in lungs of patients with COPD and alpha-1 antitrypsin deficiency (AAT), but not fibrotic lung diseases, suggesting that this activated EC state is a hallmark of degenerative emphysematous lung disease. This comprehensive and long-term atlas of tissue regeneration in the lung demonstrates the enormous cellular plasticity in its response to injury, enumerating the many transient and persistent cell states that arise after acute injury to the lung and their correlation to human lung disease.

## Results

### The proliferative response to viral injury is asynchronous across cellular compartments of the lung

Using 5-ethynyl-2’-deoxyuridine (EdU) staining, we previously observed that lung EC proliferation peaks during the third week after influenza injury, substantially later than epithelial proliferation^9,28^. To track proliferation of facultative progenitors as they are activated after injury in an unbiased way, we bred *Ki67^ires-CreERT2^* mice with *ROSA26^LSL-tdTomato^*mice, hereafter referred to as *Ki67^CreERT2^; ROSA26^LSL-tdTm^* mice^29,30^. These mice were divided into five cohorts that received tamoxifen at different time points to induce lineage tracing in proliferating cells and their progeny. The first group received tamoxifen at homeostasis, and uninjured lungs were collected seven days after administration. The other four cohorts were infected with H1N1 PR8/GP33 influenza A virus (IAV). Each cohort received two doses of tamoxifen at a different time after IAV infection to label proliferating cells (**Fig. 1A**). We sacrificed mice with adequate weight loss from each cohort the week following tamoxifen administration and used expression of the fluorescent tdTomato (tdTm) protein to quantify the number of proliferating cells and their descendants across each time window (**Fig. S1A**). We gated proliferating cells into cellular compartments using flow cytometry for tdTm^+^ Ki67-traced cells in combination with surface markers for immune cells (CD45), ECs (CD31), epithelial cells (CD326; EPCAM), and mesenchymal cells (lineage-negative) (**Fig. S1B-C**). Immune proliferation peaked during the 2-7 dpi time window and the proportion of Ki67-traced immune cells remained relatively high over the ensuing three weeks, likely representing the influx and persistence of bone marrow-derived immune cells in the lung following infection (**Fig. 1B, C**). Mesenchymal proliferation peaked during the 7-15 dpi time window and waned in the third week post infection (**Fig. 1B, C**). Epithelial proliferation also peaked in the 7-15 dpi time window and was sustained into the third week post infection, whereas endothelial proliferation continued to increase and peaked during the 14-22 dpi time window (**Fig. 1B, C**). By 21-28 dpi, proliferation in all lineages had returned to levels observed in uninjured mice (**Fig. 1B, C**). The same patterns of proliferation were observed using immunofluorescence (IF) and RNAscope analysis of Ki67-tracing in CD45^+^ immune cells, *Pdgfra*^+^ and *Pdgfrb*^+^ mesenchymal cells, NKX2-1^+^ epithelial cells, and ERG^+^ ECs (**Fig. 1D-G, Fig. S1D-G**). These data demonstrate that proliferation occurs asynchronously across the course of lung regeneration and suggests that signaling from compartments that respond early, such as immune cells, shape the regenerative outcomes in compartments that respond later.

**Figure 1.**
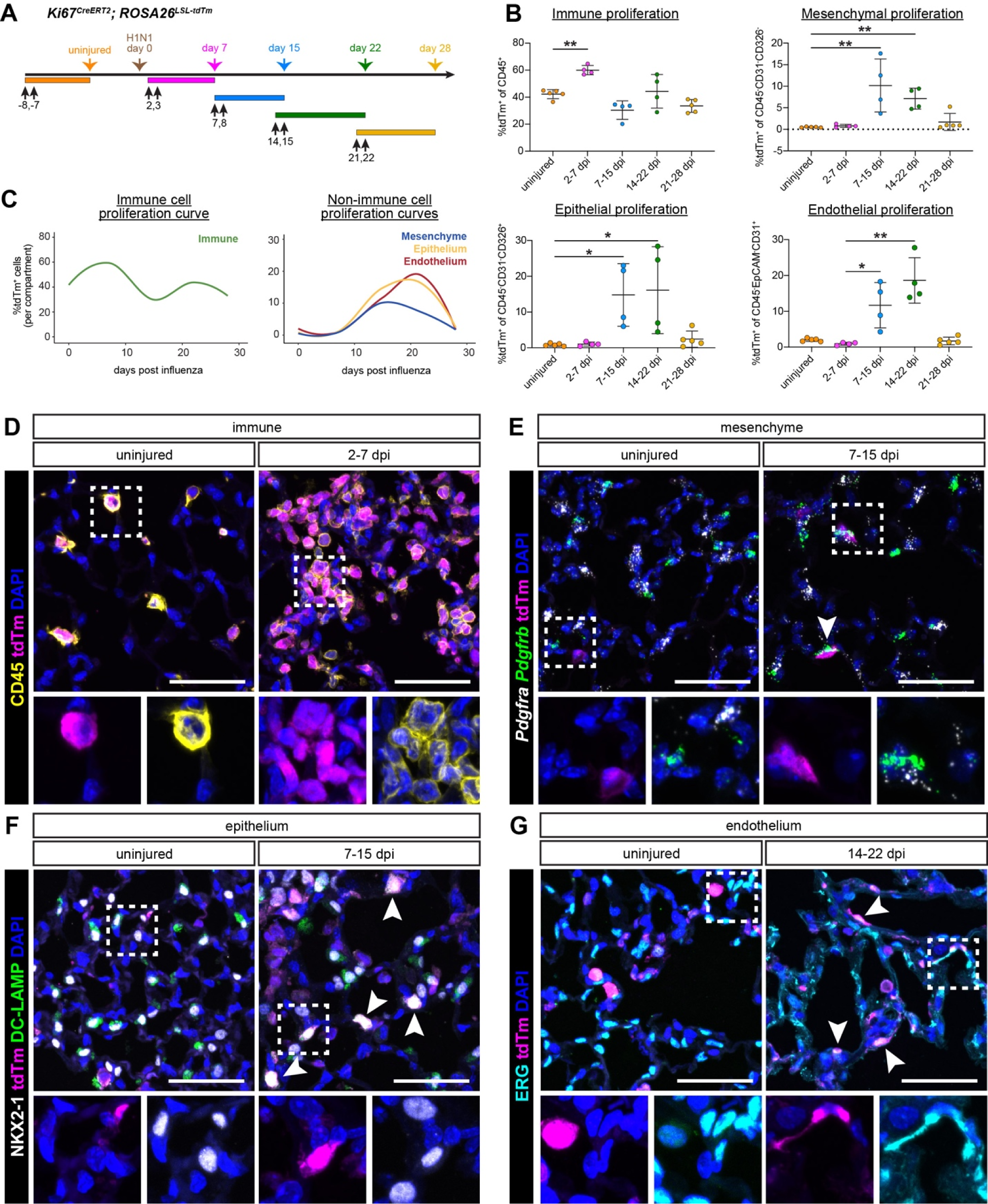
Proliferation in response to viral infection is asynchronous across compartments of the lung. **(A)** Schematic of experimental setup for flow cytometry analysis of *Ki67^CreERT2^; ROSA26^LSL-tdTm^* mice. Black arrows indicate time points for tamoxifen administration; colored arrows indicate sample collection for flow cytometry. **(B)** Changes in proliferation over the response to injury, indicating flow cytometric detection of %tdTm^+^ cells in each compartment during each time window. For all comparisons, a Kruskal-Wallis test was performed followed by Dunn’s multiple comparisons test. * p<0.05, ** p<0.01, *** p <0.001. **(C)** Local polynomial regression fitting (LOESS) in R was used to generate proliferation curves from the flow cytometry data quantified in (B). Samples collected from uninjured mice are represented as being collected on ‘Day 0’. **(D-G)** Immunofluorescence (IF) for the *Ki67^CreERT2^; ROSA26^LSL-tdTm^* lineage trace (tdTm, magenta) in combination with: **(D)** IF for immune cells (CD45^+^) at homeostasis and 2-7 dpi, **(E)** RNAscope for mesenchymal cells (Pdgfra^+^ and/or Pdgfrb^+^) at homeostasis and 7-15 dpi, **(F)** IF for AT2 epithelial cells (NKX2-1^+^/DC-LAMP^+^) at homeostasis and 7-15 dpi, **(G)** IF endothelial cells (ERG^+^) at homeostasis and 14-22 dpi. Scale bars 50 μm.

### The regenerating lung is transcriptionally and proliferatively heterogeneous

To better understand how cellular responses vary through time, we generated a single-cell transcriptional atlas of lung regeneration by performing scRNA-seq of digested whole lungs from lineage traced *Ki67^CreERT2^; ROSA26^LSL-tdTm^* mice at various time points post-IAV infection (**Fig. 2A**). To assess the extent of injury in uninfected versus infected lungs, we collected one lung from each mouse for hematoxylin and eosin (H&E) staining and used our previously described computational lung damage assessment program to bin lung tissue regions into three zones corresponding to severely damaged, damaged, and normal tissue structure^17^. As expected, the percentage of severely damaged lung tissue increased over time following IAV infection and peaked at 19 dpi, with areas remaining at 42 dpi (**Fig. S2A,B**). scRNA-seq data from all 25 animals were merged to create a dataset composed of 123,189 cells representing immune, endothelial, epithelial, and mesenchymal compartments identified based on expression of canonical markers (**Fig. S2C,D**). We subset the dataset by regenerative time point and observed similar cellular contributions to each compartment from each collection day (**Fig. S2C,E**). We further annotated each compartment’s cellular constituents using expression of cell type-specific marker genes defined by the LungMap Consortium and observed the expected cell types in each compartment^31^ (**Fig. 2B,C**).

**Figure 2.**
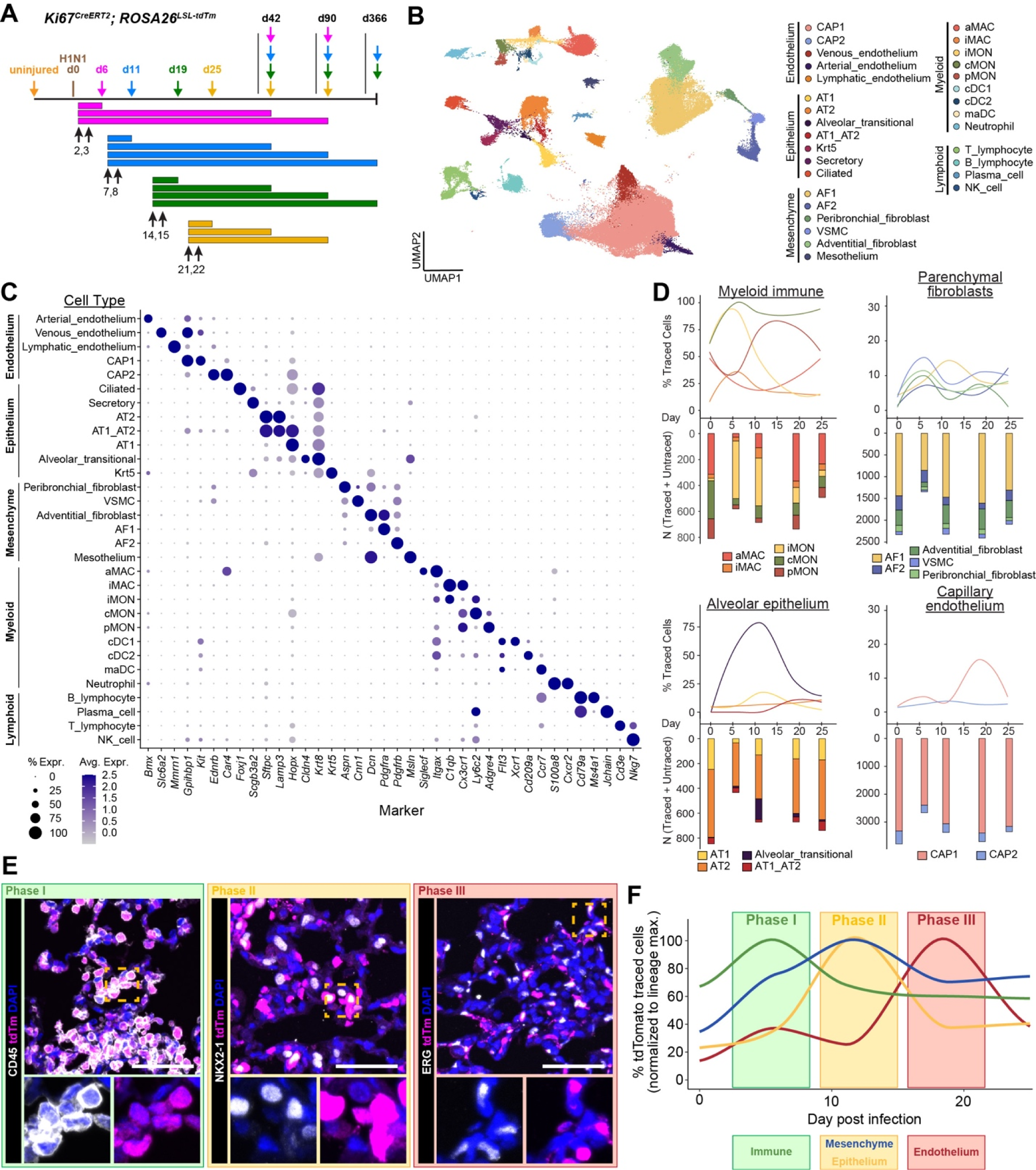
Mapping transcriptional and proliferative heterogeneity in the regenerating lung. **(A)** Schematic of experimental setup for scRNA-seq analysis. Black arrows indicate tamoxifen administration; colored arrows indicate sample collection for scRNA-seq and histology. One cohort of mice (n=2) received two doses of tamoxifen at homeostasis, and uninjured lungs were collected at 3- or 42-days post tamoxifen. Each experimental cohort received tamoxifen to induce lineage labeling at a different time after IAV infection: 2-3 dpi, 7-8 dpi, 14-15 dpi, or 21-22 dpi. Lungs were collected from each cohort at either 72-96 hours post-tamoxifen induction (6, 11, 19, and 25 dpi, respectively; n=2 per cohort), 42 dpi (n=2 per cohort), 90 dpi (n=1 per cohort), or 366 dpi (n = 3 total) **(B)** Merged scRNA-seq data from the 25 mice shown in (A), following quality filtering and annotation. **(C)** Expression of marker genes in each annotated cell type. **(D)** Percentage of informatically detected Ki67-traced cells and cell counts for samples in uninjured mice and each short time window (3-6, 8-11, 15-19, and 22-25 dpi). ‘Untraced’ cells and cells from later time points (42, 90, and 366 dpi) were excluded from this analysis. Samples collected from uninjured mice are represented as being collected on ‘Day 0’. **(E)** Representative images of Ki67-traced immune cells (CD45^+^tdTm^+^) during Phase I, 3-6 dpi; traced epithelial cells (NKX2-1^+^/tdTm^+^) during Phase II, 8-11 dpi; and traced endothelial cells (ERG^+^/tdTm^+^) during Phase III, 15-19 dpi. Scale bars 50 μm. **(F)** Lineage level proliferation curves include all cell types shown in (B) and (C). Data for each lineage are depicted as a proportion of the maximum value for that lineage.

To identify lineage-traced proliferating cells and their descendants, we informatically detected cells expressing tdTomato from the *ROSA26^LSL-tdTomato^*locus (**Fig. S2F**) and compared compartment- and cell type-specific proliferative dynamics. Focusing on the subset of samples collected 3-4 days after Cre-mediated recombination allowed us to resolve the kinetics of recently proliferating cells. We quantified Ki67-tracing in the most common cell types in the alveolar lung parenchyma, including myeloid immune cells such as monocytes and macrophages, parenchymal fibroblasts and smooth muscle cells, alveolar epithelial cells, and capillary endothelial cells. We observed comparatively high levels of Ki67-tracing in cells of the monocyte-macrophage lineages (**Fig. 2D**). In the mesenchymal compartment, *Pdgfra*^+^ alveolar fibroblast 1 (AF1) cells exhibited a modest proliferative response, while *Pdgfrb*^+^ AF2 cells were largely untraced (**Fig. 2D**). Within the alveolar epithelial compartment, we observed that alveolar transitional epithelial cells were by far the most traced cell type, corresponding to their dramatic accumulation at 11 dpi and disappearance shortly thereafter (**Fig. 2D**). While this method showed a lower level of AT2 proliferation than the alveolar transitional epithelial cells, this likely reflects a potent and rapid entrance of proliferative AT2 cells into the alveolar transitional cell state. Within the capillary endothelium, CAP1 ECs exhibited robust proliferative tracing, while CAP2 ECs were largely untraced (**Fig. 2D**).

IF analysis of proliferating NKX2-1^+^ epithelial cells, ERG^+^ endothelial cells, and CD45^+^ immune cells in tissue sections of mice used for scRNA-seq confirmed these findings (**Fig. 2E, Fig. S2G**). The summation of these data, combined with our EdU and flow cytometry data^9^, reveals that lung regeneration occurs in three distinct phases characterized by separate peaks in proliferation for immune, epithelial, mesenchymal, and endothelial cells (**Fig. 2F**).

### Alveolar macrophages are reconstituted bidirectionally from lung-resident and bone marrow-derived cells through an inflammatory intermediate

Recent work has demonstrated that acute lung injury leads to both the apoptosis of alveolar macrophages (aMACs)^32^ and the infiltration of a mixed population of myeloid immune cells, including neutrophils and inflammatory monocytes (iMONs)^33^. Over the subsequent weeks, iMONs differentiate into cells resembling alveolar macrophages, although they exhibit a range of phenotypic differences compared to native aMACs, such as lower expression of the surface protein SiglecF, higher expression of interleukin 6 (IL-6) and Tnf, and increased chromatin accessibility at the *C1q* locus^34–37^. Over the ensuing months after infection, iMONs take on the surface phenotype of lung-resident aMACs and participate in surfactant recycling^36^. The aMAC compartment is also partially reconstituted from surviving macrophages that proliferate *in situ*^38^. Given these potential origins for aMACs, we used our lung regeneration atlas to explore the range of intermediate states available and transcriptional choices made during aMAC differentiation.

The longitudinal lung regeneration atlas data was subset on the myeloid immune compartment, identifying canonical resident immune cell populations of the lung including aMACs, interstitial macrophages (iMACs), conventional and plasmacytoid dendritic cells, and neutrophils (**Fig. 3A**). Three major clusters of monocytes were observed: classical monocytes (cMONs) were defined by high expression of *Ly6c2* and low expression of *Cx3cr1*, while cells with low expression of *Ly6c2* and high expression of *Cx3cr1* were defined as non-classical or patrolling monocytes (pMONs). A third population of monocytes expressed intermediate levels of *Ly6c2,* accumulated during the first two weeks after infection, and was present mainly during infection and early regeneration (**Fig. 2C**, **Fig. 3A,B**). These were defined as inflammatory monocytes (iMONs) given their close temporal association with inflammation. During Phase I (2-6 dpi), corresponding to the peak of immune infiltration of the lung^33^ and the height of immune proliferation in our dataset, we observed a dramatic loss of aMACs and expansion of iMONs (**Fig. 3A,B**). Over the subsequent three weeks, the aMAC compartment was progressively reconstituted from two seemingly distinct cellular sources: a group that transcriptionally resembled aMACs present at homeostasis (**Fig. 3B, asterisk**), and a group passing along a trajectory originating from iMONs present at 6 dpi (**Fig. 3B**). By 90 dpi, the aMAC compartment returned to a state resembling homeostasis (**Fig. 3B**), which is consistent with observations from bleomycin injury^36^.

**Figure 3.**
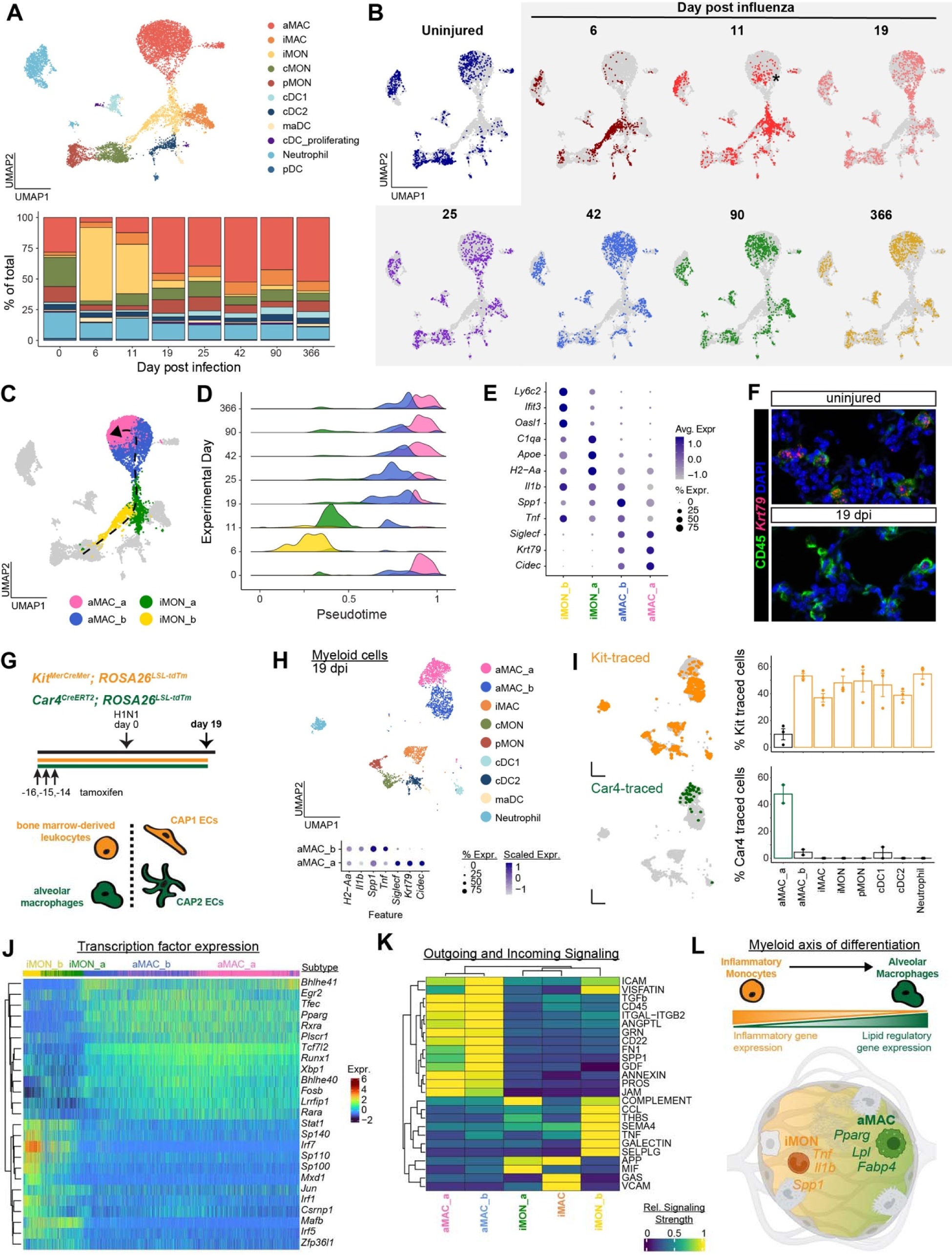
Differentiation of inflammatory monocytes to alveolar macrophages proceeds through a transient inflammatory cell state. **(A)** Myeloid immune cells in the scRNA dataset were re-clustered, re-projected in UMAP space, and summarized proportionally by experimental day. **(B)** Myeloid immune cells were evenly down sampled by experimental day and highlighted in the re-projected UMAP space. **(C)** aMAC and iMON cells were re-clustered and projected with new assignments onto the same UMAP space (cells of other lineages shown in grey). Dashed line reflects the pseudotime axis of differentiation. **(D)** aMAC and iMON subtypes summarized in ridge plot reflecting compositional changes along the pseudotime axis over experimental time. **(E)** Marker gene identification was performed and selected genes reflecting the inflammatory signatures of iMON and aMAC cells are shown. **(F)** Tissue sections from uninjured mice or IAV infected mice at 19 dpi were stained for CD45 and *Krt79*, demonstrating decreased *Krt79* staining after infection. **(G)** Schematic depicting the *Kit^MerCreMer^* and *Car4^CreERT2^* lineage tracing experiment. **(H)** Myeloid immune cells from the *Kit^MerCreMer^*and *Car4^CreERT2^* lineage trace experiment were re-clustered and projected in UMAP space. The aMAC compartment is color coded by subtype while other cell types are rendered according to the color scheme in (A). A subset of aMAC marker genes shown in (E) are depicted. **(I)** Bioinformatic detection of traced cells from *Kit^MerCreMer^; ROSA26^LSL-tdTm^* and *Car4^CreERT2^; ROSA26^LSL-tdTm^* mice. **(J)** Differential expression of transcription factors over pseudotime was evaluated using tradeSeq. The top 25 differentially expressed TFs are represented, ordered from left to right along the pseudotime axis. **(K)** Ligand-receptor signaling in myeloid immune cells. Rows representing signaling pathways and columns labeling cellular subtypes were arranged using unsupervised k-means clustering. **(L)** Schematic demonstrating differentiation of iMON cells into aMACs. Abbreviations: aMAC, alveolar macrophage; iMAC, interstitial macrophage; iMON, inflammatory monocyte; cMON, classical monocyte; pMON, patrolling (non-classical) monocyte; iMON, inflammatory monocyte; cDC1, conventional dendritic cell 1; cDC2, conventional dendritic cell 2; maDC, mature dendritic cell; pDC, plasmacytoid dendritic cell.

Cells in the iMON and aMAC lineages were reclustered at a higher resolution, resulting in two clusters of iMONs and two main clusters of aMACs. Using supervised pseudotime analysis, we generated an axis of differentiation originating with the ‘iMON_b’ cluster, which expanded dramatically at 6 dpi, and terminating with the ‘aMAC_a’ cluster representing mature alveolar macrophages, present at highest numbers in uninjured mice and samples from 90 and 366 dpi (**Fig. 3C, D**). This trajectory suggested a course of differentiation that progressed from iMON_b to iMON_a, aMAC_b, and finally aMAC_a, mirroring the temporal distribution (**Fig. 3D**). Several groups have reported macrophage populations that emerge during acute injury and express high levels of inflammatory cytokines such as Spp1, IL-1β and Tnf^33,39–41^. Our findings were consistent with these observations: at 6 dpi the predominant iMON_b population expressed high levels of transcripts encoding interferon response proteins such as *Ifit3* and *Oasl1* (**Fig. 3E**). By 11 dpi the iMON population no longer expressed high levels of interferon response genes but had a gene signature closer to interstitial macrophages (iMACs), including expression of *C1qa, Apoe,* and *H2-Aa* (**Fig. 3E**). This iMON_a state appeared to be transcriptionally intermediate to aMACs, iMACs, and dendritic cells, consistent with the reported contribution of recruited monocytes to each of these lineages^36,42–44^.

Since the contribution of monocytes to interstitial macrophages is well established^45,46^, we generated a second pseudotime trajectory originating with the iMON_b cluster and ending with iMACs (**Fig. S3A**). To determine the transcriptional programs most characteristic of the iMON-aMAC and iMON-iMAC trajectories, we performed gene set enrichment analysis using the genes whose transcription varied most over each axis of differentiation. Lipid regulatory programs were among the most variable gene sets for the iMON-aMAC trajectory, consistent with upregulation of genes involved in surfactant phospholipid catabolism such as *Pparg* (**Fig. S3B**)^34,47^. In contrast, the iMON-iMAC axis was defined by transcriptional changes in genes involved in host responses to pathogens, such as upregulation of genes encoding complement proteins as well as MHC-II (**Fig. S3B**).

At homeostasis, most aMACs belonged to the aMAC_a cluster. In contrast, during regeneration the aMAC_b cluster predominated, with a gradual return to the pre-injury composition by 90 dpi (**Fig. 3D**). Transcriptionally, the aMAC_b cluster exhibited upregulation of several inflammatory cytokines, as well as downregulation of *Siglecf* as previously reported^33,36^. In contrast, the aMAC_a cluster expressed higher levels of the canonical aMAC marker *Krt79* as well as *Cidec* (**Fig. 3E**)^48^. At homeostasis, we observed large numbers of macrophages expressing surface CD45 and intracellular *Krt79* RNA (**Fig. 3F**). In contrast, at 19 dpi, most CD45^+^ cells that morphologically resembled macrophages expressed low or undetectable levels of *Krt79* (**Fig. 3F**).

To directly evaluate whether the aMAC_b cluster originated from lung resident aMACs polarized *in situ* or from recruited iMONs, we sought to specifically label either lung resident aMACs or bone marrow-derived myeloid cells. We therefore crossed *ROSA26^LSL-tdTm^* mice with a previously published Kit^MerCreMer^ line^49^ and a Car4^CreERT2^ line we generated to study the capillary endothelium (**Fig. 2C, 3G; Fig. S3C**). scRNA-seq was performed at 19 dpi to examine the contribution of both *Car4*^+^ and *Kit*^+^ cells to the myeloid compartment (**Fig. S3D-F**). Sub-clustering the myeloid fraction of this scRNAseq data revealed two transcriptionally distinct populations of aMACs corresponding to the aMAC_a and aMAC_b clusters we had identified previously (**Fig. 3H**). The aMAC_a cluster consisted of cells that were lineage traced in the Car4^CreERT2^ mice, while the aMAC_b cluster consisted of cells traced by Kit (**Fig. 3I**). These data demonstrate that during lung regeneration, the alveolar macrophage compartment is reconstituted from both bone marrow-derived iMONs and surviving aMACs. iMON-derived aMACs have a transient ‘inflammatory’ transcriptional signature that persists throughout active regeneration, while surviving aMACs and their traced progeny transcriptionally resemble the aMACs present at homeostasis. By 90 dpi, however, the inflammatory signature that initially characterizes bone marrow-derived aMACs has subsided, and the aMAC pool uniformly resembles the population present at homeostasis.

To evaluate the differences in cell identity and state over the axis of differentiation, we analyzed expression of transcription factors (TFs) within the Animal Transcription Factor Database^50^. As expected, the iMON_a cluster that dominated at 6 dpi expressed high levels of interferon responsive TFs including *Irf1, Irf5,* and *Irf7* (**Fig. 3J**). In contrast, the fully differentiated aMAC_a cluster expressed high levels of *Pparg* and its heterodimer binding partner *Rxra*^51^, as well as *Bhlhe40* and *Bhlhe41*^52^. *Mafb*, which is required for the differentiation of iMONs into iMACs, was downregulated over the iMON-aMAC axis of differentiation, consistent with its dispensability for aMACs (**Fig. 3J**)^53^. To better understand differences in cell signaling that accompany the adaptation of aMAC cells to the alveolar niche, we used the CellChat analytic suite^54^ to perform ligand-receptor analysis on the cells within the monocyte-macrophage subset of our data. Unsupervised clustering of the pooled ‘outgoing’ and ‘incoming’ signaling pathway activities demonstrated that cells belonging to the aMAC_a and aMAC_b subtypes shared a relatively similar signaling profile that was strikingly different from iMACs and both iMON clusters. Within the aMAC lineage, the mature aMAC_a cluster exhibited lower expression of genes involved in several inflammatory pathways such as SPP1 and TNF (**Fig. 3K**). Our time-resolved lineage tracing data establish an axis of differentiation for cells of the monocyte-macrophage lineage wherein monocytes expressing high levels of interferon response genes are recruited to the lung and progress along an axis of differentiation, leading to downregulation of inflammatory genes. Differentiation is characterized by upregulation of TFs required for the execution of homeostatic aMAC functions, such as surfactant lipid catabolism, and the downregulation of TFs favoring the iMAC developmental trajectory (**Fig. 3L**).

### Epithelial regeneration is characterized by the development of transient and persistent cell states

Following injury, alveolar epithelial cells proliferate in response to inflammatory cues such as IL-1, Tnf, and other factors supplied by recruited monocytes^21,24,25^. Homeostasis is further restored by the clearance of oxidized surfactant lipids and reestablishment of normal alveolar surface tension, which is controlled in part by aMACs^55,56^. In the epithelium, the previously identified alveolar transitional cell state accounts for the bulk of the proliferation within the alveolar epithelium as measured by the percentage of Ki67-traced cells in our scRNA-seq dataset (**Fig. 2D**)^21–23^. We therefore sought to determine whether and how transitional cells and other conceivable injury-induced cell states contribute to the alveolar injury response, and whether the epithelium fully returns to homeostasis as regeneration proceeds.

At homeostasis we observed the expected cell types, including ciliated and secretory cells, AT1 and AT2 cells, and a population of cells expressing markers of both AT1 and AT2 cells that we designated ‘AT1_AT2’ (**Fig. S4A**). AT1_AT2 cells are unlikely doublets since their UMI counts, feature counts, and doublet detection score counts were comparable to those of alveolar transitional cells (**Fig. S4B**). Moreover, this cell state has been previously reported in human and mouse datasets^16,57,58^.

To focus on alveolar regeneration rather than the well-described dysplastic epithelial response^59,60^, we subset the alveolar epithelial compartment into AT1 and AT2 cells, alveolar transitional epithelial cells, and AT1_AT2 cells, while excluding airway cells and Krt5^+^ airway-derived cells (**Fig. 4A**). We observed that the accumulation of alveolar transitional epithelial cells peaked at 11 dpi; these cells diminished in number over time and were only rarely observed by 25 dpi and thereafter (**Fig. 4A**). Alveolar transitional cells were much more strongly traced than the other intermediate cell type, AT1_AT2 cells, suggesting that alveolar epithelial proliferation after IAV infection passes preferentially through this state (**Fig. 4B**). In contrast, AT1_AT2 cells were present in similar number across most time points and had Ki67-tracing levels consistent with AT1 and AT2s. Unlike transitional cells, AT1_AT2 cells were present in a fairly consistent proportion at all time points except 6 dpi and had a transcriptional profile intermediate to AT1 and AT2 cells with no unique marker genes (**Fig. 4A,C**). These data demonstrate that the alveolar transitional cell serves as the predominant intermediate state during inflammation and injury, while AT1_AT2 cells might represent a population of poised or partially differentiated cells at other times.

**Figure 4.**
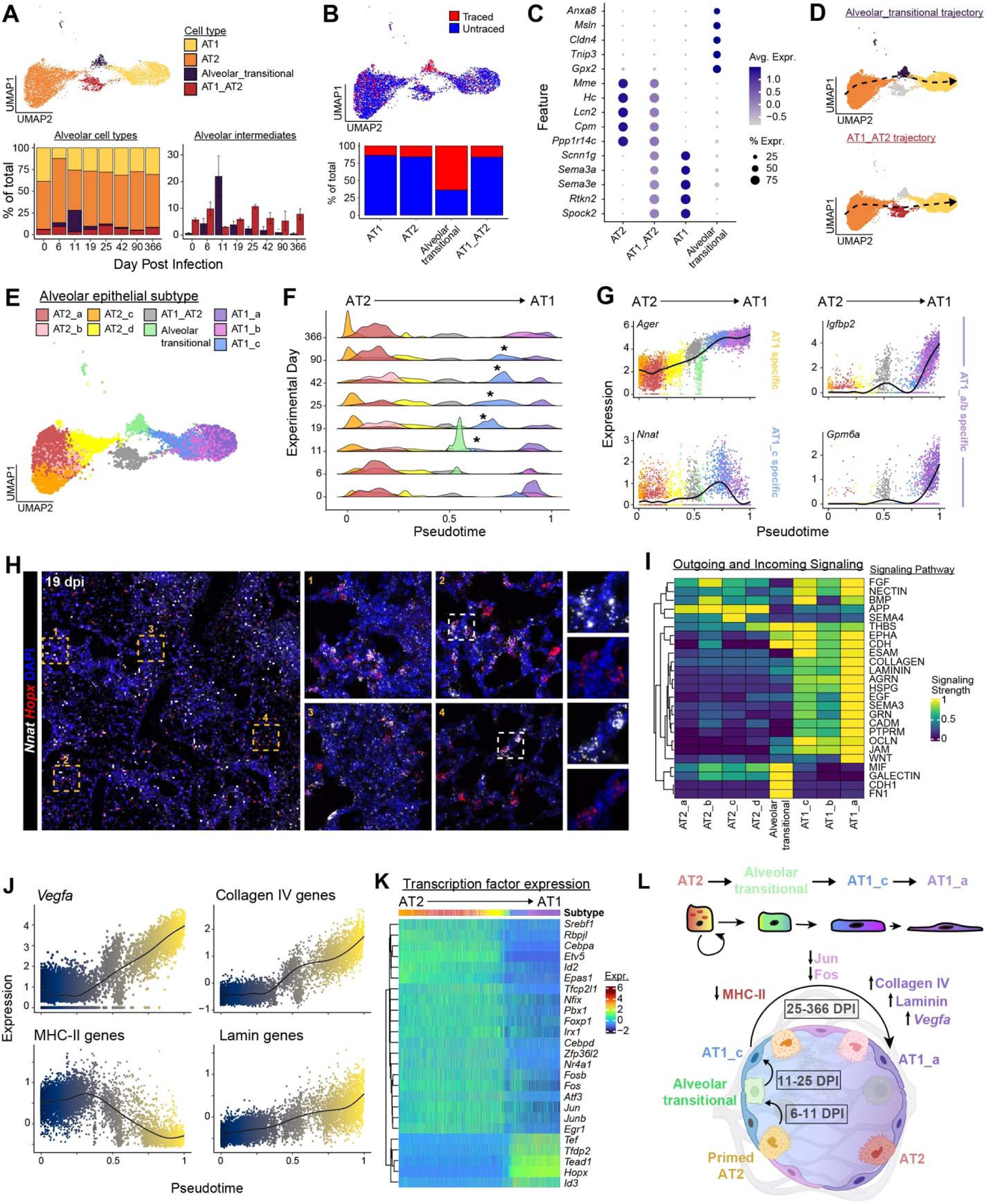
Transient and long-lasting epithelial transition states arise following IAV infection. **(A)** Alveolar epithelial cells in the scRNA dataset were re-clustered, projected in UMAP space, and summarized by experimental day**. (B)** Traced and Untraced alveolar epithelial cells plotted in UMAP space and summarized by cell type. **(C)** Dot plot for the top 5 marker genes in AT1, AT2, and alveolar transitional cells. No genes were found to be specific to AT1_AT2 cells in this analysis. **(D)** Supervised trajectory analysis was performed on subsets of the alveolar epithelial compartment. The ‘Alveolar_transitional trajectory’ and ‘AT1_AT2 trajectory’ are shown in dashed lines. Cells included in each trajectory were highlighted in each case and color coded as in (A). **(E)** Alveolar epithelial cells were re-clustered at higher resolution and projected onto the same UMAP coordinates as in (A). **(F)** Cells were ordered along a normalized pseudotime trajectory representing the average of the two trajectories shown in (D) and scaled from 0 to 1, with 0 denoting the AT2 origin. Ridgeline plots demonstrating cellular composition over experimental time are colored as in (E). **(G)** Expression of the AT1 marker gene *Ager*, along with marker genes for the AT1_c and AT1_a clusters through pseudotime. **(H)** Representative histologic sections from mice at 19 dpi, with RNAscope using *Hopx* and *Nnat*. Inset panels 1-4 show representative sections. Image brightness was adjusted in the low magnification image to improve visualization of fluorescent cells. **(I)** CellChat analysis of pathways that varied across the AT2 to AT1 differentiation axis, top 20 pathways shown. Dendrograms are clustered using unsupervised k-means clustering. **(J)** Expression of cell type defining transcriptional programs over pseudotime. For ligands encoding ECM proteins (i.e. collagen IV and laminin) or MHC, where multiple genes are expressed in a similar pattern across cell subtypes, a module score was generated to simplify data presentation using the Seurat AddModuleScore function. **(K)** Differential expression of transcription factors over pseudotime, top 25 TFs are shown. **(L)** Schematic depicting differentiation of AT2 cells to AT1_a cells via intermediate cell states.

To better understand the cellular dynamics of the alveolar epithelium during regeneration, we re-clustered the epithelium at a higher resolution and generated pseudotime trajectories originating with AT2 cells and terminating with AT1 cells (**Fig. 4D,E**). Projecting all alveolar epithelial cells along this axis revealed a population of AT1 cells that began accumulating at 11 dpi, persisted through 90 dpi, and was no longer distinct by 366 dpi (**Fig. 4F, asterisk; Fig. S4C,D**). This AT1_c state exhibited substantially higher Ki67-tracing at 11dpi but not at later time points, suggesting that they derive from alveolar transitional cells (**Fig. S4E**). Cells in the AT1_c cluster had similar expression of the AT1 marker gene *Ager* as other AT1 clusters but expressed increased levels of *Nnat,* an imprinted gene that plays a role in brain development and metabolism (**Fig. 4G**)^61,62^. RNAscope analysis demonstrated the accumulation of a *Hopx^+^* population of AT1 cells with high *Nnat* expression at 19 dpi, consistent with the expansion of AT1_c cells (**Fig. 4H**). In contrast, we observed a separate cluster of ‘AT1_a’ cells that was present in uninjured mice and at all time points after injury, marked by the expression of *Igfbp2* and *Gpm6a* (**Fig. 4G**). *Igfbp2* has been previously identified as a marker of terminally differentiated AT1 cells^63^, supporting the hypothesis that AT1_c cells represent an immature state that terminally differentiates over the course of regeneration.

To elucidate changes in signaling within the alveolar niche that vary due to the regeneration process, we performed ligand-receptor analysis across the AT2-to-AT1 differentiation axis. AT1_c cells appeared intermediate to AT1_a and AT2 cells, further supporting the possibility that they represented an incompletely differentiated cell state (**Fig. 4I**). To test this hypothesis, we focused on core features of AT2 and AT1 biology such as expression of ECM proteins and Vegfa in AT1 cells^5,64^ and MHC-II in AT2 cells^65^. Feature and module scores indicating expression of genes associated with mature AT1 cell functions, such as *Vegfa* and the ECM proteins collagen IV and laminin, increased across the axis of differentiation from AT2 cells to AT1_a cells (**Fig. 4J**). In contrast, expression of MHC II genes decreased across the trajectory (**Fig. 4J**). In each case, expression of the given gene or gene set was intermediate in AT1_c cells (**Fig. 4J**). We next examined expression changes in TFs along the axis of alveolar epithelial differentiation. We observed the expected decrease in expression of AT2-specific transcription factors such as *Etv5*^66^, *Cebpa*^67^, and *Tfcp2l1*^16^, along with an increase in AT1-specific transcription factors such as *Hopx* and *Tead1*^68^ (**Fig. 4K, S4F**). Other transcription factors such as *Fos* and *Jun* were expressed at higher levels in AT1_c compared to AT1_a cells (**Fig. 4K, S4F**). These data suggest that incomplete differentiation of AT1 cells and the presence of immature AT1_c cells at 90 dpi could have functional consequences for AT1 cell function following injury (**Fig. 4L**).

### The pulmonary endothelium is highly sensitive to interferon signaling

The phased nature of the proliferative response to IAV indicates that the temporal sequence of signaling between compartments may determine how each cell type in the lung mounts a regenerative response. The pulmonary endothelium is unique in its late proliferative response (**Fig. 2D-F**), suggesting that ECs either require pro-proliferative signals from other regenerating compartments, or receive signals that dampen proliferation early in regeneration. One hallmark of the early response to IAV is immune cell secretion of interferons (IFNs) and inflammatory cytokines, and the pulmonary epithelium is known to respond strongly to inflammatory signaling^16,17,24,25^. We therefore created an “interferon score” for each cell in our scRNA-seq dataset based on expression of 87 IFN-stimulated genes (ISGs)^69^ (**Fig. 5A,B**). Interferon scores varied substantially across time following infection, with the highest level of ISG expression occurring at 6 dpi in each compartment. Remarkably, the endothelium had the highest interferon score of all non-immune compartments (**Fig. 5B**). Examination of differentially expressed genes for CAP1s at 6 dpi revealed that viral and interferon responses were consistently enriched (**Fig. S5A**). IF analysis for ERG in combination with nuclear pSTAT1, a marker of active interferon signaling, confirmed that pulmonary ECs are actively responding to IFN signaling at 6 dpi but not at homeostasis or 19 dpi (**Fig. 5C, Fig. S5B**).

**Figure 5.**
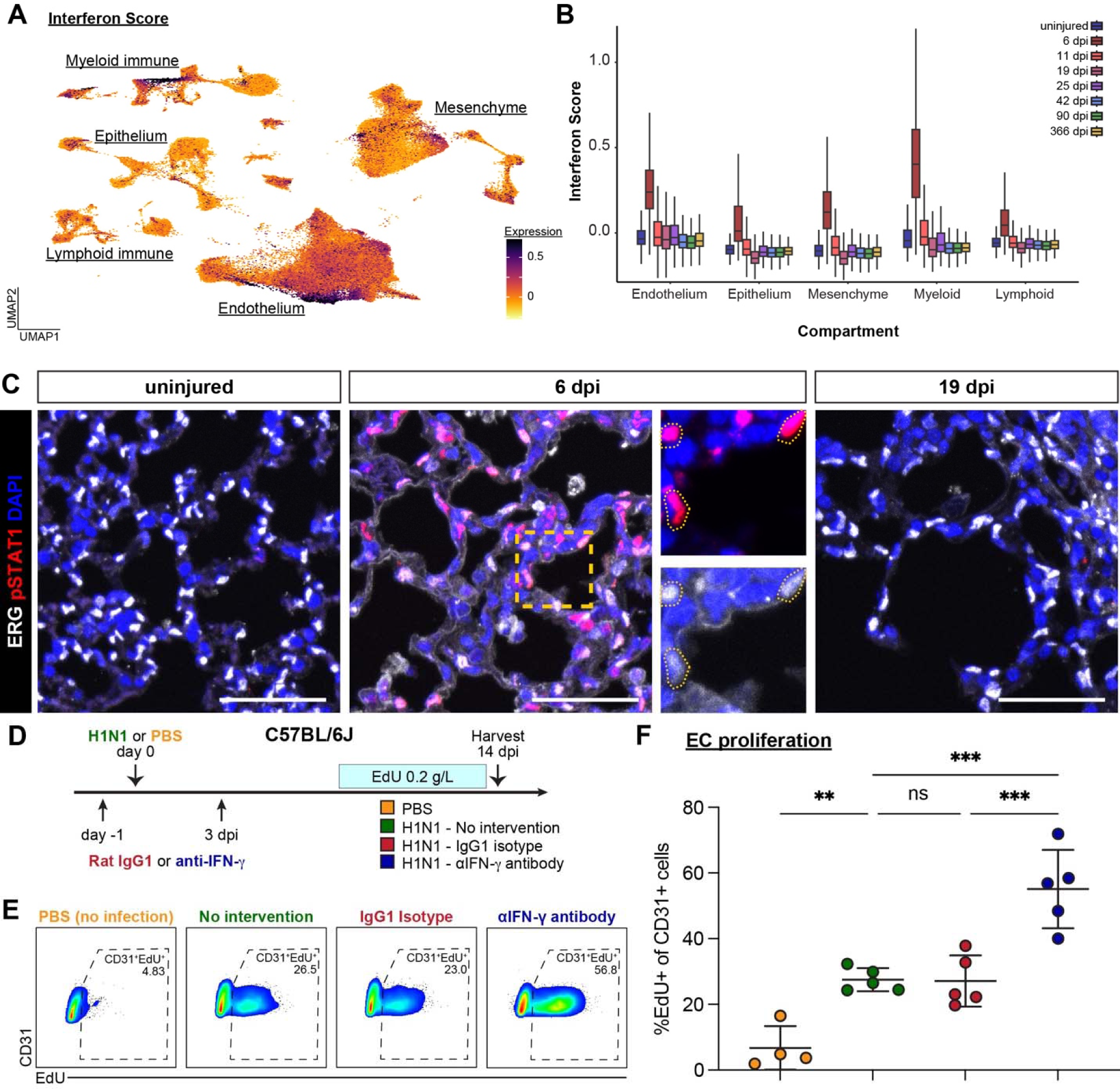
The pulmonary endothelium responds strongly to interferon signaling. **(A)** Interferon module score for cells in the Ki67 lineage trace experiment. Darker colors indicate higher expression of interferon stimulated genes. **(B)** Quantification of interferon module score in lineages and over the experiment. **(C)** IF for the nuclear EC marker ERG as well as pSTAT1. Scale bars 50 μm. **(D)** Schematic the IFN-γ blocking experiment. All mice received EdU in their drinking water from 7-14 dpi. **(E)** Representative flow cytometry plots for EdU incorporation in CD31^+^ ECs. **(F)** Quantification of flow cytometry in (E). One-way ANOVA, p<0.0001, followed by Holm-Sidak’s multiple comparisons test. **, p< 0.01. ***, p<0.001.

Interferon signaling is known to repress cell proliferation to slow viral replication^70–72^. Examination of interferon receptor expression demonstrated near-ubiquitous expression of the IFN-α receptors *Ifnar1* and *Ifnar2* but high expression of IFN-γ receptor 1 (*Ifngr1*) in ECs, particularly in CAP1s (**Fig. S5C,D**). To determine whether high levels of IFN signaling in ECs is responsible for their late proliferative response, we blocked IFN-γ signaling using an IFN-γ blocking antibody or rat IgG1 isotype control and examined EC proliferation (**Fig. 5D**)^73^. Mice that received the IFN-γ blocking antibody or isotype control had equivalent weight loss and percentage of CD45^+^ cells in lung homogenates at 14 dpi compared to IAV-infected mice with no antibody treatment (**Fig. S5E**). In contrast, mice treated with anti-IFN-γ had approximately two-fold higher EC proliferation compared to IAV-infected mice receiving isotype or no intervention (**Fig. 5E,F**). This indicates that IFN-γ signaling is one factor responsible late EC proliferation during regeneration.

### A population of injury-induced capillary ECs (iCAPs) arises after injury and persists indefinitely

Recent studies have demonstrated the existence of EC heterogeneity in the context of lung injury and repair^7,9,19,11,74^. To determine whether injury-associated endothelial states arise after influenza and whether they persist, we subset the longitudinal scRNA-seq atlas data to examine only capillary ECs (**Fig. 6A**). We observed previously identified capillary EC subtypes, including CAP1s, which proliferate after lung injury to restore the capillary EC pool^7,9^, and CAP2s, which are regulated by Vegf signaling from the epithelium and have been suggested to play an integral role in gas exchange^7,9,64^. We also observed a distinct group of ECs intermediate to the CAP1 and CAP2 transcriptional states that was not present at homeostasis, emerged around 11 dpi, and persisted through 366 dpi (**Fig. 6A,B**). We termed this EC state “injury-induced capillary endothelial cells,” or iCAPs. iCAPs were characterized by high expression of the CAP1 markers *Gpihbp1* and *Kit*, low expression of the CAP2 markers *Ednrb* and *Car4*, expression of several unique marker genes including *Sparcl1* and *Ntrk2*, and upregulation of MHC-II and *Ifngr1* (**Fig. 6C,D**).

**Figure 6.**
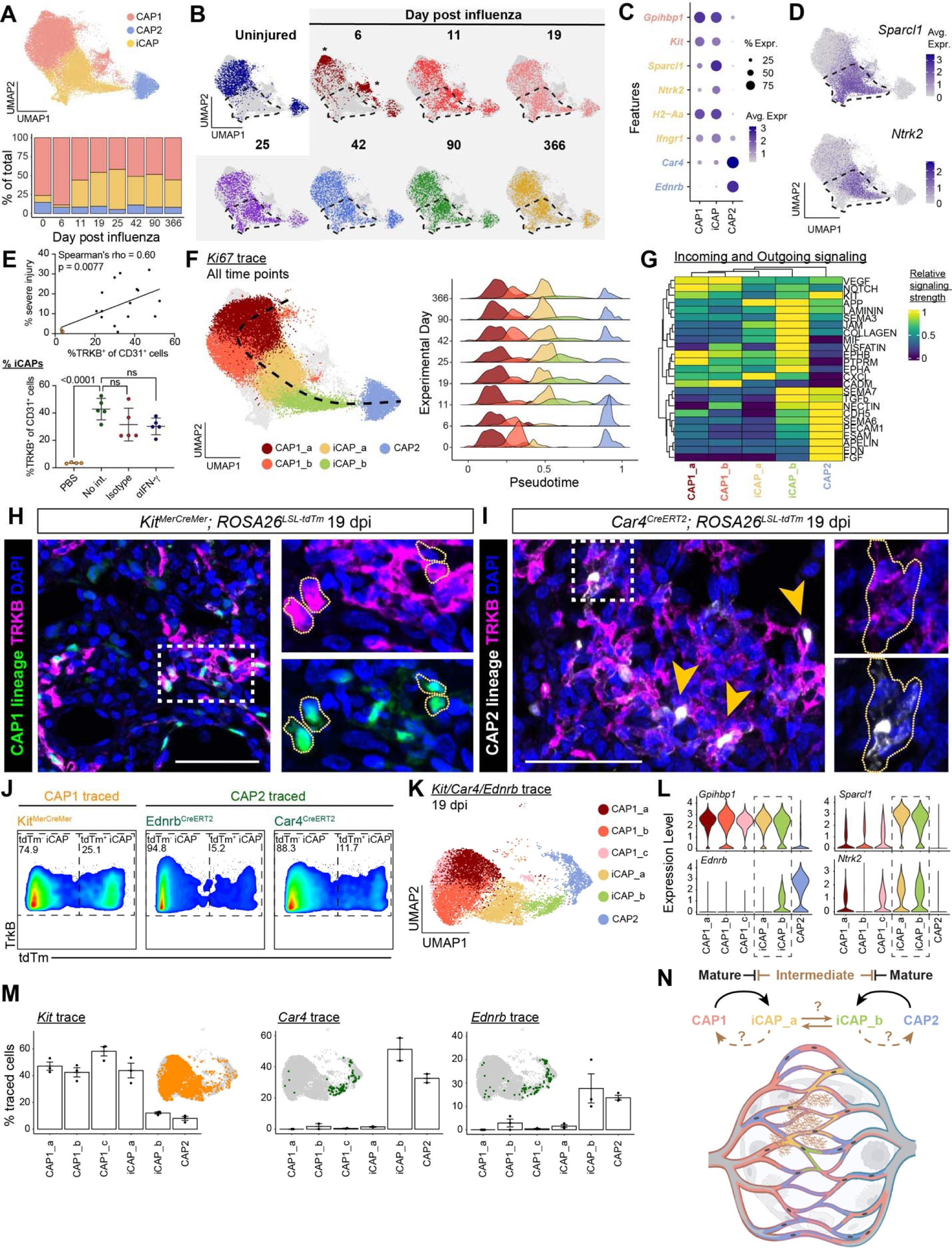
An injury-induced capillary endothelial cell subtype arises at 11 days post injury. **(A)** Capillary endothelial cells in the scRNA dataset were re-clustered, projected in UMAP space, and summarized by experimental day. **(B)** Re-clustered cells in the capillary endothelial compartment were evenly down sampled by experimental day and highlighted accordingly. Dashed lines denote the approximate boundaries of the iCAP cell state. Asterisks denote capillary EC clusters that are mainly present at 6 dpi. **(C)** Expression of genes specific to CAP1 ECs (*Gpihbp1*, *Kit*), CAP2 ECs (*Car4*, *Ednrb*), and iCAP ECs (*Sparcl1*, *Ntrk2*, *H2-Aa, Ifngr1*) in capillary ECs. **(D)** Expression of iCAP-specific markers genes. **(E)** (Top) Injury score calculation was performed from mice in the IFN-γ blocking experiment. The percent ‘severe’ histologic injury was correlated with the percent CD31^+^TRKB^+^ iCAP cells identified using flow cytometry. Mice from the mock infected group are colored in orange. (Bottom) % CD31^+^TRKB^+^ cells in each group. One-way ANOVA, p<0.0001, followed by Holm-Sidak’s multiple comparisons test. **(F)** (Left panel) Cells in the capillary EC compartment were reclustered at higher resolution. A pseudotime trajectory was generated originating in the CAP1_a subtype and ending with CAP2 cells. Cells from excluded EC subtypes (CAP1_c, CAP1_d, iCAP_c) are shown in grey. (Right panel) The distribution of EC subtypes along the pseudotime axis is highlighted over experimental time using ridge plots. **(G)** CellChat analysis of outgoing and incoming signaling pathways in capillary endothelial cell groups. **(H)** IF for the iCAP marker TrkB (the protein product of *Ntrk2*) in CAP1 lineage traced mice (*Kit^MerCreMer^; ROSA26^LSL-tdTm^*). **(I)** IF for TrkB in CAP2 lineage traced mice (*Car4^CreERT2^; ROSA26^LSL-tdTm^*). **(J)** Representative flow cytometry plots demonstrating tdTm tracing of iCAPs in CAP1 and CAP2 lineage traced ECs (gated on CD45^−^CD31^+^TRKB^+^ cells). **(K)** The capillary endothelium was reclustered from *Kit^MerCreMer^*(CAP1), *Ednrb^CreERT2^* and *Car4^CreERT2^* (CAP2) lineage traced mice at 19 dpi were subset as in (F). The same subtypes of capillary ECs were identified apart from an additional CAP1_c cluster. **(L)** Expression of the canonical CAP1 and CAP2 markers *Gpihbp1* and *Ednrb*, as well as the iCAP state markers *Sparcl1* and *Ntrk2* in capillary subtypes from the data shown in (K). Dashed box highlights iCAP subtypes. **(M)** Bioinformatic detection of tdTm expression in capillary EC subtypes. **(N)** Schematic demonstrating known and hypothetical relationships between canonical capillary EC subtypes and iCAP subtypes.

Endothelial *Sparcl1* expression has recently been reported to exacerbate viral pneumonia in mice^75^. RNAscope analysis for *Sparcl1* expression in combination with pan-EC marker *Pecam1* and CAP1 marker *Gpihbp1* identified *Sparcl1^+^*CAP1s at 11 dpi (**Fig. S6A,B**), although *Sparcl1* staining was not specific to the endothelial compartment. Sparcl1^+^ ECs were found exclusively in damaged areas of the lung following influenza, providing further evidence that the iCAP state is induced by injury and may depend on the damage-associated niche for its emergence (**Fig. S6C**). Flow cytometry data from the IFN-γ blocking experiment further demonstrated that the proportion of iCAPs expressing TrkB (the protein product of *Ntrk2*) correlated with histologic injury severity (**Fig. 6E, Fig. S6D,E**). Additionally, TrkB^+^ ECs were rarely detected in uninfected mice, although the percentage of TrkB^+^ cells did not vary significantly in anti-IFN-γ treated mice (**Fig. 6E**).

Since CAP1s are the main proliferative EC type following viral lung injury (**Fig. 2D**) and can differentiate into CAP2 cells^7,64^, we hypothesized that the iCAP state might represent a differentiation intermediate between CAP1s and CAP2s. Ki67-tracing of iCAP cells was intermediate to CAP1 and CAP2 cells, suggesting that they could represent an admixture of CAP1- and CAP2-derived cells polarized toward a common transcriptional intermediate (**Fig. S6F**).To further resolve the cellular heterogeneity within the capillary endothelium, ECs were reclustered at a higher resolution, revealing four CAP1 clusters, three iCAP subtypes, and a single CAP2 group (**Fig. S6G**). To simplify our analysis, we focused on the most numerous capillary EC clusters (CAP1_a, CAP1_b, iCAP_a, iCAP_b, CAP2), while excluding two CAP1 clusters that were predominantly present at 6 dpi (CAP1_c, CAP1_d; **Fig. 6B, asterisk**) and the iCAP_c cluster which did not express *Ntrk2* (**Fig. S6G,H**). Unlike the myeloid and epithelial intermediate states, the iCAP_a and iCAP_b clusters persisted for at least one year after injury (**Fig. 6A,F**). Ligand-receptor interaction analysis within the capillary endothelium demonstrated that iCAP_a cells were most similar to CAP1 cells, while the iCAP_b and CAP2 cells diverged functionally from CAP1s (**Fig. 6G**). This finding also supported the possibility that the iCAP state was composed of a mixture of CAP1- and CAP2-derived cells.

### The iCAP state arises through plasticity of both CAP1 and CAP2 cells

To define the cell of origin for the two iCAP subtypes, we analyzed scRNA-seq data from the Kit^MerCreMer^ and Car4^CreERT2^ lineage traced mice, in combination with scRNA-seq data generated from Ednrb^CreERT2^ mice that also lineage trace CAP2s (**Fig. 3G, S3C**).

IF analysis for tdTm^+^ (lineage-traced) cells and the iCAP markers Sparcl1 and TrkB was used to determine whether one or both capillary EC types could give rise to the iCAP state following injury. Kit-traced, Sparcl1-positive ECs were not observed at homeostasis in uninjured mice (**Fig. S6I**). At 19 dpi, we observed Kit-traced TrkB^+^ or Sparcl1^+^ iCAP ECs, indicating that the iCAP state is at least partially derived from CAP1 ECs (**Fig. 6H; Fig. S6I**). Ednrb-traced or Car4-traced, Sparcl1^+^ ECs were likewise not observed at homeostasis (**Fig. S6J**). Surprisingly, we also observed TrkB^+^ and Sparcl1^+^ iCAP cells bearing the Car4 and Ednrb lineage trace by IF at 19 dpi, indicating that the iCAP state is also derived in part from the CAP2 lineage (**Fig. 6I; Fig. S6J-L**). Quantification of lineage-traced ECs by flow cytometry revealed that iCAP ECs arise from the CAP1 and CAP2 lineages (**Fig. 6J, Fig. S6E**).

Comparison of the merged scRNA-seq data from Kit^MerCreMer^, Car4^CreERT2^, and Ednrb^CreERT2^ lineage-traced mice to the longitudinal time course revealed a similar profile of EC clusters, including the two dominant CAP1 subtypes, two distinct iCAP clusters, and CAP2 cells (**Fig. 6K**). As above, we observed that both iCAP clusters expressed high levels of *Sparcl1* and *Ntrk2* (**Fig. 6L**). Bioinformatic analysis of the lineage trace from these animals revealed that Kit-traced ECs were found at high levels in the CAP1 and iCAP_a subsets (**Fig. 6M**). In contrast, Car4-traced and Ednrb-traced ECs were observed in the CAP2 and iCAP_b subsets (**Fig. 6M**). Taken together, our observations by IF, flow cytometry, and scRNA-seq indicate that the iCAP cell state arises bidirectionally from CAP1 and CAP2 cells after viral lung injury, localizes to areas of significant tissue injury, and persists for at least a year after IAV infection (**Fig. 6N**).

### iCAPs represent a developmental EC state associated with human degenerative lung disease

As the iCAP state arose from both canonical EC types and did not specifically depend on IFN-γ signaling, we explored whether iCAPs might result from a disruption of the alveolar niche following injury. One major contributor to the disrupted niche is the invasion of airway derived Krt5^+^ epithelial cells that form dysplastic epithelial scar tissue^60,76,77^. Although TrkB^+^ ECs could be found in close association with Krt5^+^ epithelial cells in some areas of damaged tissue, they were also present at high numbers in severely damaged areas that did not contain Krt5^+^ epithelium but were denuded of AT1 epithelial cells (**Fig. 7A**). We also observed TRKB^+^ iCAPs in areas of damaged but regenerating tissue adjacent to areas of severe damage^17,18^, but we did not observe TRKB^+^ iCAPs in areas of normal tissue architecture (**Fig. 7A**). These data led us to conclude that the iCAP state is not strictly dependent on the presence of Krt5^+^ epithelium.

**Figure 7.**
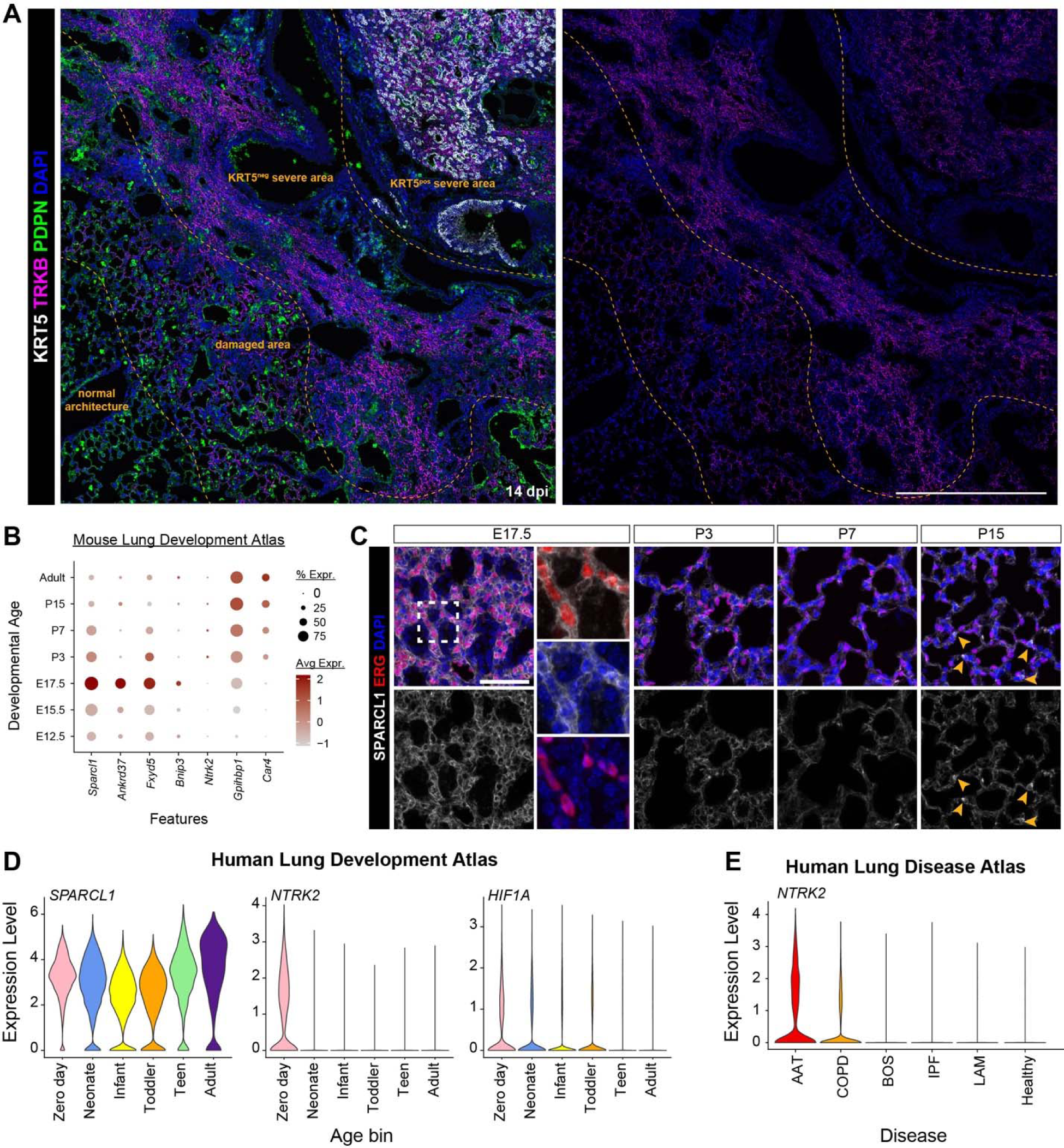
The iCAP signature is associated with development and emerges during chronic tissue injury. **(A)** Low magnification section of mouse lung at 14 dpi demonstrating a heterogeneous pattern of injury that includes severely damaged areas with and without investment of Krt5^+^ cells. PDPN^+^ cells are found in areas with normal architecture or, to a lesser extent, in damaged areas without consolidation. The right panel highlights TRKB staining alone. Scale bar, 500 μm. Image brightness was adjusted in these low magnification images to improve visualization of fluorescent cells. **(B)** Combined mouse capillary endothelial cells from a published mouse developmental atlas^6^ were binned and expression of iCAP signature genes is depicted according to developmental age bin. **(C)** IF of Sparcl1 and Erg in samples used to generate the mouse developmental atlas corresponding to (B). Scale bar 50 μm. **(D)** Violin plots of the iCAP marker genes *SPARCL1* and *NTRK2*, as well as the upstream regulator *HIF1A* in a human pediatric developmental cohort. **(E)** Expression of *NTRK2* in pooled capillary endothelial cells (includes both CAP1 and CAP2) in a human disease cohort. AAT: alpha-1 antitrypsin deficiency; COPD: chronic obstructive pulmonary disease; BOS: bronchiolitis obliterans syndrome; IPF: idiopathic pulmonary fibrosis; LAM: lymphangioleiomyomatosis.

We next examined whether iCAPs lost their lung-specific phenotype due to injury. Comparing expression of the iCAP transcriptional program to transcriptomes of ECs from other organs in an organ-specific EC atlas revealed that expression of several iCAP-specific genes was high in non-lung ECs (**Fig. S7A**)^78^. Applying an iCAP “score” comprised of the top 50 iCAP-specific genes to ECs from each organ demonstrated that ECs of other organs were more “iCAP-like” than homeostatic lung ECs (**Fig. S7B**).

To examine whether the iCAP state resembled immature lung ECs, we compared their transcriptional program to a previously published scRNA-seq dataset spanning mouse embryonic and postnatal lung development^6^. Several iCAP-specific genes, including *Sparcl1*, *Ankrd37*, *Fxyd5*, and *Bnip3*, were most highly expressed in the E17.5 embryonic mouse lung, prior to the transition to air breathing, while expression of adult CAP1 and CAP2 marker genes *Gpihbp1* and *Car4* did not increase until postnatal time points as others have observed (**Fig. 7B**)^5^. IF for SPARCL1 in embryonic and postnatal mouse lung tissue demonstrated a decrease in SPARCL1^+^/ERG^+^ ECs from E17.5 through P7, the start of alveologenesis in the mouse lung (**Fig. 7C**). To determine if this observation holds true in human lung, we examined a human lung development atlas containing scRNA-seq from human lung tissue obtained starting at the day of birth through adulthood. While SPARCL1 is not a useful iCAP marker in humans given its broad expression throughout the lifespan, *NTRK2* and its upstream regulator *HIF1A* were expressed at high levels only in the zero-day lung tissue (**Fig. 7D**). These data demonstrate that the iCAP state shares characteristics with immature human and mouse pulmonary ECs.

To determine whether the iCAP state was observed in human lung disease, a human lung disease atlas from the Penn-CHOP Human Lung Tissue Bank (HTLB) was used to compare expression of the iCAP marker *NTRK2* across different human disease states. As in the mouse, *NTRK2* expression was very low in healthy control lung tissue, but we observed high *NTRK2* expression in emphysematous diseases including COPD and alpha-1-antitrypsin deficiency (AAT) (**Fig. 7E**). In contrast, although *Ntrk2*^+^ cells have been observed in mouse lungs injured with bleomycin^79^, we did not observe expression of *NTRK2* in idiopathic pulmonary fibrosis (IPF) (**Fig. 7E**). These data implicate the iCAP transcriptional state in several human lung diseases. The presence of the iCAP state is therefore indicative of a damaged alveolar niche and may represent a tipping of the balance towards dysplastic rather than euplastic regeneration.

## Discussion

Using a multimodal approach incorporating novel genetic tools, lineage tracing, and unbiased scRNA-seq time course experiments, we have generated a comprehensive longitudinal atlas of lung regeneration following viral injury. We demonstrate that regeneration is asynchronously phased across the different lineages of the lung and characterized by a succession of proliferative responses. During the first week after infection, a wave of immune proliferation occurs, reflecting mobilization of the inflammatory response and recruitment of myeloid cells to the lung. Epithelial and mesenchymal proliferation peaks during the second week as airway and alveolar epithelial cells repair and regenerate the gas exchange interface. Unexpectedly, endothelial proliferation peaks during the third week after injury, suggesting that structural and signaling cues supplied by the other cellular compartments may be necessary for endothelial cells to respond, and that this delay in response could have an important impact on both the rate and extent of tissue regeneration.

Our data also reveal the plasticity of cellular states that arise and resolve across the lung’s cellular compartments during regeneration. At homeostasis, the lung is quiescent, with transcriptionally distinct cell types and little proliferation. During the first week after infection, broad transcriptional changes occur, corresponding to activation of interferon responses in all cellular lineages. In the second week, we observe the emergence of transitional states in the myeloid, epithelial, and endothelial compartments. By generating a comprehensive atlas of the axes of lineage differentiation, we can now appreciate the cellular dynamics of lung regeneration.

Large numbers of inflammatory monocytes are recruited to the lung during the first week after infection, coincident with the loss of resident alveolar macrophages. The aMAC compartment is then reconstituted bidirectionally from the *in situ* proliferation of Car4^+^ mature aMACs as well as the differentiation of bone marrow-derived iMONs that are lineage-traced by Kit. As iMONs traverse this axis of differentiation, they downregulate inflammatory transcriptional programs and begin expressing the lipid regulatory machinery required to reestablish homeostasis through their criticial role in recycling surfactant.

As expected, our data reveal that the alveolar epithelium is canonically regenerated by proliferation and differentiation of AT2 cells. However, the relationship between this potent proliferative response and previously described or novel transition states had remained unclear. This longitudinal atlas of lung regernation shows that following proliferation, AT2 cells are rapidly and preferentially shunted into the alveolar transitional cell state, before ultimately differentiating into AT1 cells. Although this initial differentiation is rapid, a population of *Nnat* expressing AT1 cells persists for months after infection. These immature AT1 cells express lower levels of genes involved in hallmark AT1 function, such as Vegfa and ECM protein synthesis, as well as higher levels of MHC-II. By one year after infection, this subpopulation of AT1 cells is no longer present and the AT1 compartment is similar to homeostasis. This delay in full AT1 differentiation could be a limiting factor in successful alveolar regeneration and function.

Unexpectedly, we also observe a population of transcriptionally distinct ECs, iCAPs, that emerges during the second week after influenza infection, expresses high levels of Sparcl1 and TrkB, localizes to sites of lung injury, and persists indefinitely. To better understand the cellular origins of the iCAP state, we developed a novel capillary endothelial lineage tracing approach. Using newly generated *Car4^CreERT2^* and *Ednrb^CreERT2^* mouse lines, we demonstrate that iCAP cells are transcriptionally heterogeneous and derive from both CAP1 and CAP2 cells. Finally, we asked what contextual or signaling cues led to the development of iCAP cells. While IFN-γ blockade led to precocious endothelial proliferation, it did not significantly affect the abundance of iCAP cells. To assess whether the iCAP state represents a less differentiated capillary cell state, interrogation of a mouse lung developmental data set revealed that the iCAP transcriptional program was elevated late during human and mouse prenatal lung development and subsequently downregulated early in postnatal life. We also noted that the iCAP signature was upregulated in humans with degenerative lung diseases such as COPD and AAT, suggesting that endothelial pathology may be a central feature in these conditions and providing an important association between acute injury and chronic human lung diseases^80–82^.

Our data highlight the immense complexity of the lung’s response to injury and reveal a striking cellular plasticity during alveolar regeneration. The persistence of injury-associated cellular states for months or longer suggests that tissue remodeling may occur well after the initial phases of injury and regeneration. The cellular and molecular cues that promote development of the cell states described remain largely unknown. The signaling pathways and cellular interactions that are responsible for iCAP formation and sustenance in injured areas will be important areas of research to determine if iCAPs can be reprogrammed into either of the canonical CAP1 or CAP2 EC states. It will also be important to understand if the iCAP state shares epigenetic features with ECs during development or in other organs. Such work will shed critical light on important and unexplored facets of endothelial biology and lung regeneration.

## Materials and Methods

### Human data generation

Lungs were obtained from patients undergoing lung transplantation (for disease) or at the time of organ harvest for ‘healthy’ controls. For adults a roughly 3 cm x 2 cm section of distal lung tissue was obtained, pleura and visible blood vessels were dissected away, and sections were minced into 2 mm pieces and processed into single cell suspensions as previously described^58^. For adult lungs, CD45^+^ and CD45^−^ cells were magnetically fractionated as described below. For pediatric samples, tissue was obtained from at the time of lobar resection for other conditions and normal lung was isolated remote from the region of interest. The samples were processed in a manner similar to adult tissue. The 10X Genomics Chromium Controller platform with Chromium Next GEM Single Cell 3’ Reagent Kits v3.1 was used to assemble 3’ gene expression libraries. Sequencing was performed on an Illumina Novaseq 6000 instrument.

The ‘healthy’ control tissues used in this study were from de-identified, non-used lungs donated for organ transplantation through the Prospective Registry of Outcomes in Patients Electing Lung Transplantation (PROPEL) study, approved by the University of Pennsylvania (Penn) Institutional Review Board (IRB). Informed consent was provided by next of kin or health care proxy in accordance with NIH and institutional policies. Diseased tissue was obtained from participants enrolled in PROPEL at Penn^83^. This study was approved by the Penn (IRB), and written informed consent was obtained from all diseased patients prior to enrollment. All patient identifying information was removed prior to use. While this does not meet current NIH definition for human subject research, the reported experiments followed all guidelines, regulations, and institutional procedures required for human subject research.

#### Animals

All animal procedures used to generate this data were approved by the Institutional Animal Care and Use Committee of the University of Pennsylvania. Mice were housed in groups of two to five animals per cage when possible; if singly housed, mice were supplied with extra enrichment. The following mice were obtained from the Jackson Laboratories: *Ki67^ires-CreERT2^* (Mki67^tm2.1(cre/ERT2)Cle^, strain #029803) ^29^ and ROSA26*^LSL-tdTomato^* (Gt(ROSA)26Sor^tm14(CAG-tdTomato)Hze^, strain #007914)^30^. The *Kit^MerCreMer^*mice were a kind gift from Dr. Jeffery Molkentin and are also available at the Jackson Laboratories (Kit^tm2.1(cre/Esr1*)Jmol^, strain #032052)^49^. C57BL/6J mice were also obtained from the Jackson Laboratories (strain #000664).

### Creation of Car4^CreERT2^ and Ednrb^CreERT2^ mouse lines

Using CRISPR-based homology-directed repair in mouse embryonic stem cells, a tamoxifen-inducible CreERT2 sequence was inserted at the three-prime end of the Car4 or Ednrb coding sequence after a T2A cleavage sequence (replacing the stop codon). This sequence is followed by a 3xSTOP sequence and endogenous pA/3’ regulatory elements. An FRT-flanked Neomycin resistance cassette was used for selection of positive clones during ES cell targeting. Flp-mediated recombination using the ROSA26^FlpO^ mouse allele (JAX strain #012930) was used to remove the FRT-flanked Neo cassette. This allows expression of CreERT2 under the control of the Car4 or Ednrb promoter without disturbing expression of the endogenous gene. Detection of Car4^CreERT2^ by PCR was performed as follows:

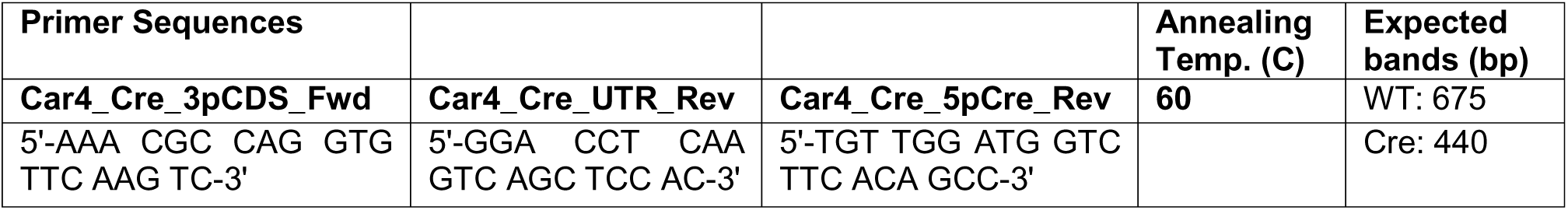

Confirmation of Neo cassette removal in the Car4^CreERT2^ line was performed as follows:

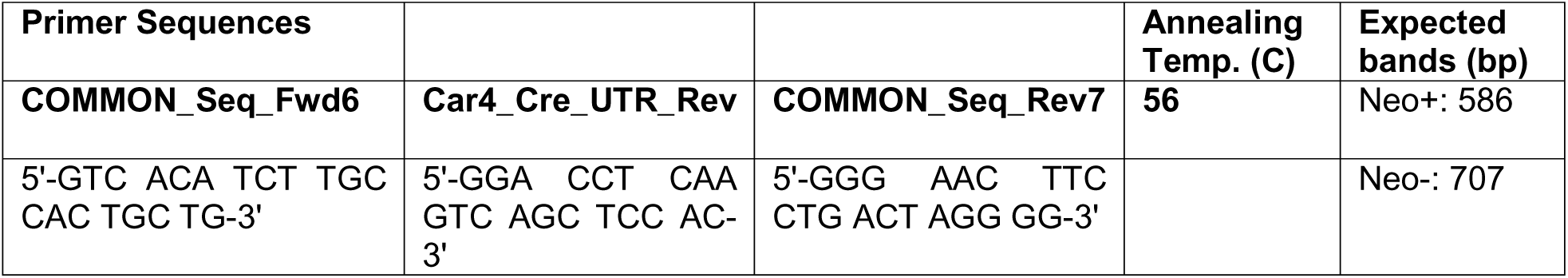

Detection of Ednrb^CreERT2^ by PCR was performed as follows:

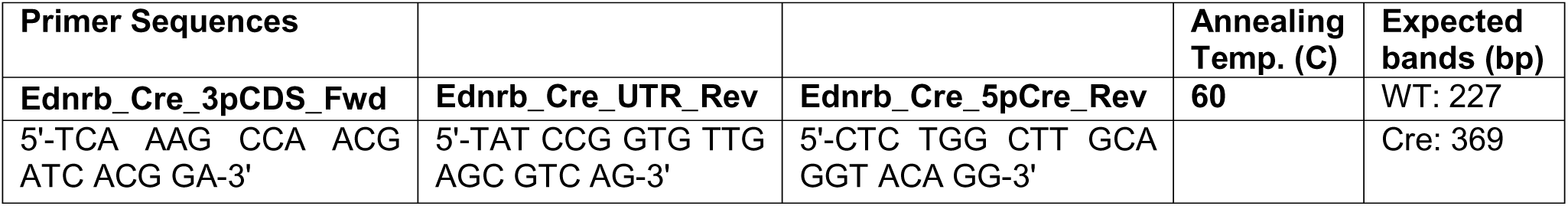

Confirmation of Neo cassette removal in the Ednrb^CreERT2^ line was performed as follows:

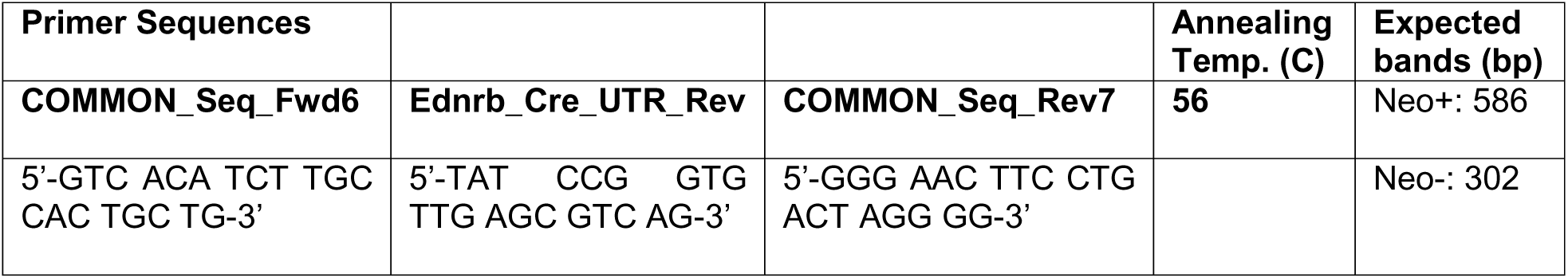

DNA construct information is available upon request. An optimal dose of tamoxifen for each line resulted in efficiencies of 50.6% and 50.1% for Car4^CreERT2^ and Ednrb^CreERT2^, respectively, and specificities of 74.6% and 91.2%, respectively.

#### Tamoxifen administration

Tamoxifen (Sigma-Aldrich #T5648) was suspended in a 1:10 mixture of ethanol and corn oil to prepare a stock concentration of 20 mg/mL. Tamoxifen solution was delivered to mice by oral gavage of 10 μL per gram body weight at the concentrations indicated. *Ki67^ires-CreERT2^*: 2 doses of 200 mg/kg; *Kit^MCM^* and *Car4^CreERT2^*: 3 doses of 200 mg/kg; *Ednrb^CreERT2^*: 1 dose of 50 mg/kg.

#### Influenza injury

The PR8-GP33 H1N1 influenza virus used to infect all genetically targeted mice was a kind gift of Dr. E. John Wherry. Mice were given a titrated dose of 1 LD_50_, determined empirically in our laboratory to be the dose at which 50% of C57BL/6J mice succumb to the infection). A commercial A/Puerto Rico/8/34 H1N1 influenza virus (ATCC #VR-95PQ, Lot 70022722) was used to infect C57BL/6J mice at a dose determined empirically in our laboratory to cause 15-20% weight loss without death (1:500,000). In all cases, H1N1 was diluted in sterile saline and administered intranasally in a volume of 50 μL. This results in a moderate level of tissue damage with heterogeneous zones of injury as previously reported ^17,18^ and as described in **Fig. S2A-B**. Mice were weighed daily for two weeks following infection and euthanized if their weight loss exceeded 30% of their original body weight. For each study, exclusion criteria were pre-established: for experiments conducted during the first week following infection, mice that loss less than 5% body weight were excluded, while for experiments conducted after the first week, mice that lost less than 15% body weight were excluded (**Fig. S1A**). If possible, each experiment contains data from at least two independent influenza infection replicates to ensure consistent results.

#### EdU administration

5-Ethynyl-2’-deoxyuridine (Santa Cruz Biotechnology, #sc-284628B, Lot C1521) was dissolved in tap water at a concentration of 2 g/L, sterile filtered, and stored at 4°C. To create a drinking water solution, this stock solution was diluted 1:10 in tap water and sterile filtered for a final concentration of 0.2 g/L. Mice consumed EdU water *ad libitum* over the time course indicated. EdU incorporation during this period was detected using the Click-iT™ Plus EdU Alexa Fluor™ 488 Flow Cytometry Assay Kit (Thermo Fisher #C10632) according to the manufacturer’s protocol.

In vivo *interferon-gamma blocking*. To block IFN-γ signaling, mice received intraperitoneal (i.p.) injections of either purified *in vivo* GOLD rat IgG1 isotype (control; Leinco Technologies #I-1195, clone GL113) or purified *in vivo* GOLD anti-mouse IFNγ (experimental; Leinco Technologies #I-1119, clone XMG1.2) one day prior to and 3 days after IAV infection. Each injection contained 1 mg of either isotype or antibody. As with all IAV infections, mice were weighed daily for two weeks following infection and euthanized if their weight loss exceeded 30% of their original body weight.

#### Histology and immunofluorescence

At the time of lung tissue harvesting, mice were euthanized with a lethal dose of CO_2_ followed by cervical dislocation. Perfusion with PBS through the right ventricle was used to clear the lungs of blood. The trachea was cannulated, and lungs were inflation-fixed with either 2% (IF) or 4% (RNAscope) paraformaldehyde in PBS, depending on the downstream application, at a pressure of 30 cm of water. Immersion fixation in 2% or 4% PFA was continued overnight at 4°C with shaking. Tissue was then dehydrated as follows: 6 x 20-minute washes in 1X PBS, overnight washes in 70% and 95% ethanol, and 2 x 1-hour washes in 100% ethanol, all at 4°C with shaking. Lung tissue was paraffin embedded and sectioned using a microtome. Hematoxylin and eosin (H&E) staining was used to assess tissue morphology and structure. Immunofluorescence analysis was used to recognize antigens using the antibodies described below. DAPI was added to slides prior to mounting. Coverslips were mounted using either Vectashield Antifade Mounting Medium (Vector Laboratories #H-1000) or Slowfade Diamond Antifade Mountant (Thermo Fisher #S36972).

| Antibody | Source | Identifiers | Concentration |
| --- | --- | --- | --- |
| CD45 (goat polyclonal) | R&D Systems | AF114, RRID:<br>AB_442146 | 1:200 |
| CXCL2 (rat monoclonal) | R&D Systems | MAB452, RRID:<br>AB_2230058 | 1:100 |
| ERG (mouse monoclonal) | Abcam | ab214341, RRID:<br>AB_3073833 | 1:100 |
| ERG (rabbit monoclonal) | Abcam | ab92513<br>RRID: AB_2630401 | 1:100 |
| LAMP3 (rat monoclonal) | Novus<br>Biologicals | DDX0191P-100<br>RRID: AB_2827532 | 1:100 |
| NKX2-1 (TTF1) (rabbit monoclonal) | Abcam | ab76013 RRID:<br>AB_1310784 | 1:50 |
| phospho-STAT1<br>(Tyr701) (58D6) (rabbit monoclonal) | Cell Signaling | 9167, RRID:<br>AB_561284 | 1:100 |
| RFP (DsRed) (rabbit polyclonal) | Rockland | 600-401-379,<br>RRID: AB_2209751 | 1:50 |
| SPARCL1 (goat polyclonal) | R&D Systems | AF2836, RRID:<br>AB_2195097 | 1:100 |
| tdTomato (goat | Origene | AB8181-200, | 1:50 |
| polyclonal) |  | RRID: AB_2722750 |  |
| TRKB (goat polyclonal) | Novus Biologicals | AF1494, RRID: AB_2155264 | 1:100 |

#### RNAscope analysis

Lung tissue was prepared as described for histology and immunofluorescence analysis above. We performed RNAscope using the Fluorescent Multiplex Reagent Kit v2 (ACD #323100) according to the manufacturer’s instructions. RNAscope probes were ordered from ACDbio and are described below. DAPI was added to slides prior to mounting. Coverslips were mounted using ProLong Gold Antifade Mountant (Thermo Fisher #P36930).

| **RNAscope probe** | **Catalog number (ACD)** |
| --- | --- |
| mm-Car4-C3 | 468421-C3 |
| mm-Gpihbp1-C3 | 540631-C3 |
| mm-Irf7-C3 | 534541-C3 |
| mm-Isg15-O1-C1 | 559271 |
| mm-Pecam1-C2 | 316721-C2 |
| mm-Sparcl1-C1 | 424641 |

#### Imaging and image analysis

All images were acquired using an LSM 710 laser scanning confocal microscope (Zeiss) at 20X and 60X. For images acquired at 60X, the Z stack function was used, and a maximal image projection was created. The tile scan function was used at 20X to acquire lower magnification images at higher resolution. Image analysis was performed in FIJI (ImageJ). Tile scans of entire lung lobes after H&E staining were acquired using the EVOS FL Auto 2 imaging system and the tile scan function at 20X. Lung damage assessment was performed using MATLAB as previously described^17^.

#### Single-cell isolation and flow cytometry

Following right ventricle perfusion of the mouse as described for histology and immunofluorescence analysis above, lung lobes were dissected from the main-stem bronchus and minced first with dissecting scissors and then with a razor blade. Minced lung tissue was digested for 35 minutes at 37°C in a combination of collagenase-I (480U/mL), dispase (100 μL/mL), and DNase (2 μL/mL) with periodic shaking, and filtered through 100 and 40 micron filters. Red blood cells were lysed using ACK lysis buffer, and single cells were resuspended in FACS buffer composed of 1% FBS in PBS. Fluorescently conjugated antibodies used for flow cytometry are described below. Cells were incubated in primary antibody for 30 minutes at 4°C (on ice) and secondary antibody, if applicable, for 30 minutes at 4°C (on ice). To analyze cell proliferation, cells were fixed in 4% PFA and permeabilized using a saponin-based reagent. Cells that had incorporated EdU were fluorescently labeled using the Click-iT™ Plus EdU Alexa Fluor™ 488 Flow Cytometry Assay Kit (Thermo Fisher #C10632) according to the manufacturer’s instructions. Flow cytometry data was acquired using an LSR Fortessa (BD Biosciences) or a Cytoflex SRT (Beckman Coulter). Analysis of flow cytometry data was performed using FlowJo software.

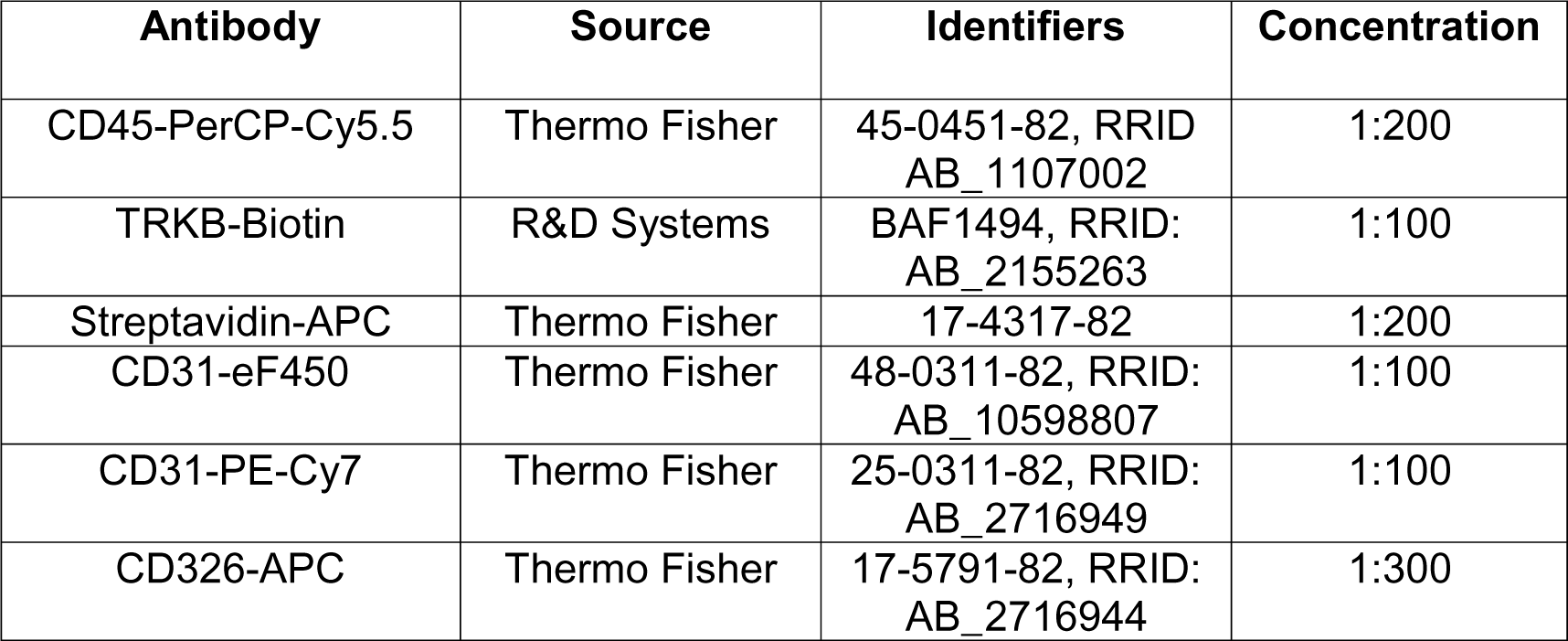

#### Magnetic-activated cell sorting

For studies involving mice, cells intended for scRNA-seq were separated into CD45-positive and CD45-negative populations using mouse CD45 MicroBeads magnetic beads (Miltenyi #130-052-301) according to the manufacturer’s instructions. Briefly, each single-cell suspension prepared as described above was spun down at 300g for 10 minutes. Each cell pellet was resuspended in 20 uL CD45 MicroBeads in 180 uL FACS buffer and incubated on ice for 30 minutes. Cells were washed twice with FACS buffer, resuspended in FACS buffer, and loaded onto two pre-washed LS columns (Miltenyi #130-042-401) per sample. A QuadroMACS separator (Miltenyi #130-091-051) was used to collect CD45-negative cells in the column flow-through. After 3 washes of the column with FACS buffer, each column was removed from the QuadroMACS separator and CD45-positive cells were eluted in 5 mL FACS buffer per column using the provided plunger. Cells were spun down and resuspended in a minimal volume (1-2 mL) of FACS buffer for counting prior to single-cell RNA sequencing.

#### Mouse single-cell RNA sequencing

To create samples for scRNA-seq, CD45-negative and CD45-positive populations for each sample acquired using MACS were combined in a proportion of 90% CD45^−^:10% CD45^+^ or 85% CD45^−^ :15% CD45^+^. Cells were counted using the Countess II FL Automated Cell Counter (Thermo Fisher). Cell suspensions were adjusted to a concentration of 800-1200 cells per microliter (8e5 - 1.2e6 cells per mL) using FACS buffer. Each sample was loaded onto a 10X Chromium Controller (10X Genomics) according to the manufacturer’s instructions. cDNA and sequencing libraries were prepared using 10X Genomics reagents and following the manufacturer’s protocol. cDNA and library quality control were performed using an Agilent 2100 Bioanalyzer using the Bioanalyzer High Sensitivity DNA kit (Agilent #5067-4626). Sequencing was performed by Azenta Biosciences using a NovaSeq instrument.

#### Mouse single-cell RNA sequencing analysis

Reads were aligned to the mouse reference genome (mm39/mGRC39) and unique molecular identifier (UMI) counts were obtained using STAR-Solo (v2.7.9a). Ambient RNA contamination was removed using SoupX v1.6.0^86^, and doublet detection were performed using scds and scrublet{Citation}. For further processing and downstream analysis, we used Seurat v4.9^87^ in R v4.3. Experiments utilizing the *Ki67^ires-CreERT2^* line were analyzed separately from those utilizing the *Kit^MCM^, Car4^CreERT2^*, and *Ednrb^CreERT2^* mice. After merging libraries from mice in each cohort, variable feature selection and annotation of reads from the ROSA26 locus was performed. Cells meeting any of the following were excluded from downstream analysis: >15% mitochondrial reads, feature count < 50, feature count > 2x median absolute deviation. The cell cycle phase prediction score was calculated using Seurat function CellCycleScoring. Data was normalized and scaled using the SCTransform function, and the effects of percent fraction of mitochondria, number of features per cell, and number of UMIs per cell were regressed out. Linear dimension reduction was done via PCA. Data was clustered using the Louvain graph-based algorithm in R. Cluster resolution was determined empirically with a resolution of 1.0 used for initial clustering of all cells and values ranging from 0.4 to 1.2 were used for analyses where reclustering within a given cellular lineage was performed. The Uniform Manifold Projection (UMAP) data reduction algorithm was used to project the cells onto two-dimensional coordinates. Clusters were then assigned to a cellular lineage based on annotation with canonical marker genes described in published references and LungMAP and elsewhere. For intra-cluster gene expression differences, the FindMarkers function was used to identify variation between specified clusters. Additional filtering was performed to remove clusters without a strong marker gene signature (which were often near the low end of the feature count cutoff), with detection of markers from multiple cellular compartments in the same cell despite passing doublet detection thresholding, and for rare cell types that were not the focus of this manuscript (i.e. platelets, erythrocytes, mast cells, and basophils).

In the Ki67-traced dataset we identified a cluster of endothelial cells expressing markers of both CAP1 and CAP2 cells with dramatically higher UMI and feature counts relative to other endothelial cell types. We removed these cells from further analysis because we could not confidently determine if they represented individual cells, doublets, or some intermediate state (e.g. binucleated or syncytial cells). As described in the main text, we also observed a cluster of cells expressing both AT1 and AT2 markers that we included in downstream analysis. We retained these cells because the feature and UMI counts were modestly elevated relative to AT1 and AT2 cells but in line with other epithelial cells (e.g. alveolar transitional and secretory cells).

For lineage tracing, cells were annotated based on expression of transcripts mapping to either the 871bp region overlapping the 3x SV40 polyA or the 1kbp region overlapping the bGH polyA following the tdTomato coding sequence. These regions which are labeled site A/B were chosen because of the higher sequencing depth. A cell was annotated as ‘Traced’ if >50% of transcripts from the ROSA26 locus were derived from the recombined allele. If no transcripts from the ROSA26 locus were present, the cell was annotated as ‘Not_detected’. The remaining cells which had detectable expression from the ROSA26 locus with <50% of transcripts originating from the recombined allele were annotated as ‘Untraced.’ The same threshold for annotation was employed for all Cre lineages used here.

Ligand receptor analysis was performed using CellChat v1.6^54^. Cell types or cellular subtypes were evenly downsampled prior to analysis. Ingoing, outgoing, and combined signaling matrices were manually retrieved from the ‘netAnalysis_signalingRole_heatmap’ function and heatmaps were generated using the ComplexHeatmap package and rendered using the ‘viridis’ package.

Gene set enrichment (GSE) and gene ontology (GO; gseGO) analysis were performed using the ‘clusterProfiler’, ‘enrichplot’, and ‘DOSE’ packages. For gseGO analyses, 10,000 permutations were performed. Heatmaps were generated using ComplexHeatmap.

Pseudotime trajectory analysis was performed using the ‘slingshot’ and ‘tradeseq’ packages^88,89^. Analyses were supervised to generate trajectories corresponding to known or hypothesized axes of differentiation. This was accomplished by removing other cell types or cellular subtypes from the data set during trajectory identification. The cells that were included are colored according to their annotation, with cells that were not included rendered in grey. The ‘fitGAM’ function in tradeseq was used to identify genes that varied along the pseudotime trajectory. Normalized pseudotime was calculated by rescaling the pseudotime coordinates from 0 to 1.

#### Graphical and statistical analysis

The ‘MetBrewer’ package in R was used to guide color palette generation for scRNA sequencing analysis. Statistical analysis of scRNA-seq data was performed in R when applicable. All other statistical analysis was performed using GraphPad Prism software. If more than two groups were analyzed, a one-way ANOVA was performed to determine if differences between groups were statistically significant, followed by Holm-Sidak’s multiple comparison tests to compare individual groups. If the conditions for parametric statistics were not met, a Kruskal-Wallace test was used to determine if differences between groups were statistically significant, followed by Dunn’s multiple comparison tests to compare individual groups. Unless otherwise indicated, all graphs display the mean and standard deviation from the mean. On all graphs, each dot shown represents a biological replicate or a single mouse. Exact p values and statistical tests used are indicated in the figure legends.

#### Data and Code availability

The scRNA-sequencing data described here has been deposited in the NCBI Gene Expression Omnibus (GEO) repository under accession number GSE262927. Code has been deposted on GitHub at (https://bit.ly/3vqeXUZ).

## Acknowledgments

The authors would like to thank Dr. E. John Wherry at Penn for providing the PR8/G33 influenza virus mouse-adapted strain. We would also like to acknowledge and thank the core facilities at Penn without whom we could not have completed this work. Flow cytometry data in this manuscript were generated in the Penn Cytomics and Cell Sorting Shared Resource Laboratory at the University of Pennsylvania (RRID: SCR_022376). This core facility is partially supported by the Abramson Cancer Center NCI grant P30 016520. Confocal microscopy was performed in the Penn Cell and Developmental Biology Microscopy Core (RRID: SCR_022373). We also thank the members of the Morrisey lab for their helpful suggestions and discussions over the course of this work.

## Author Contributions

Conception and design of the study: T.K.N., J.D.P., and E.E.M.

Acquisition, analysis, and interpretation of data: T.K.N., J.D.P., S.L., L.I.L., S.Z.

Resources: M.C.B., E.C., A.N.N., D.B.F., J.M.D.

Code, data visualization: J.D.P., M.P.M., A.B.

Writing: T.K.N., J.D.P., and E.E.M.

---

**Figure S1.**
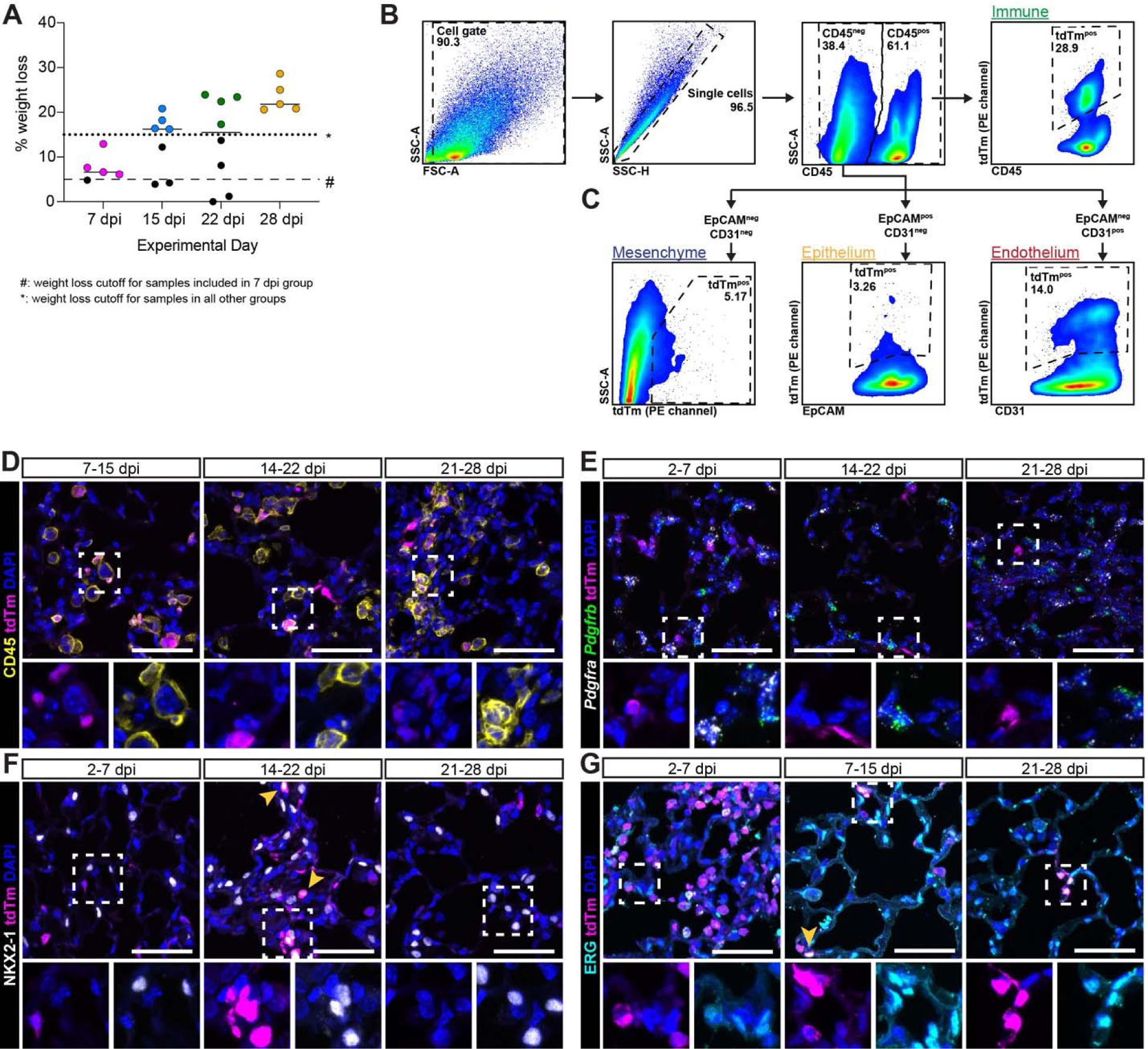
Analysis of proliferation in *Ki67^CreERT2^; ROSA26^LSL-tdTm^* animals. **(A)** Percent weight lost in animals chosen for analysis in each cohort. Black dots indicate mice that did not have adequate weight loss and were not analyzed. **(B,C)** Representative flow cytometry plots demonstrating the gating strategy used to identify Ki67-traced tdTm^+^ immune (CD45^+^), endothelial (CD45^−^EpCAM^−^CD31^+^), epithelial (CD45^−^CD31^−^EpCAM^+^), and mesenchymal (CD45^−^ CD31^−^EpCAM^−^) cells. **(D-G)** Immunofluorescence (IF) for the lineage trace (tdTm, magenta) in combination with IF or RNAscope for time points not included in Fig. 1D-G. **(D)** Immune cells (CD45^+^) at 7-15, 14-22, and 21-28 dpi. **(E)** Mesenchymal cells (*Pdgfra*^+^ and/or *Pdgfrb*^+^) at 2-7, 14-22, and 21-28 dpi. **(F)** Alveolar epithelial cells (NKX2-1^+^) at 2-7, 14-22, and 21-28 dpi. **(G)** Endothelial cells (ERG^+^) at 2-7, 7-15, and 21-28 dpi. Scale bars 50 μm.

**Figure S2.**
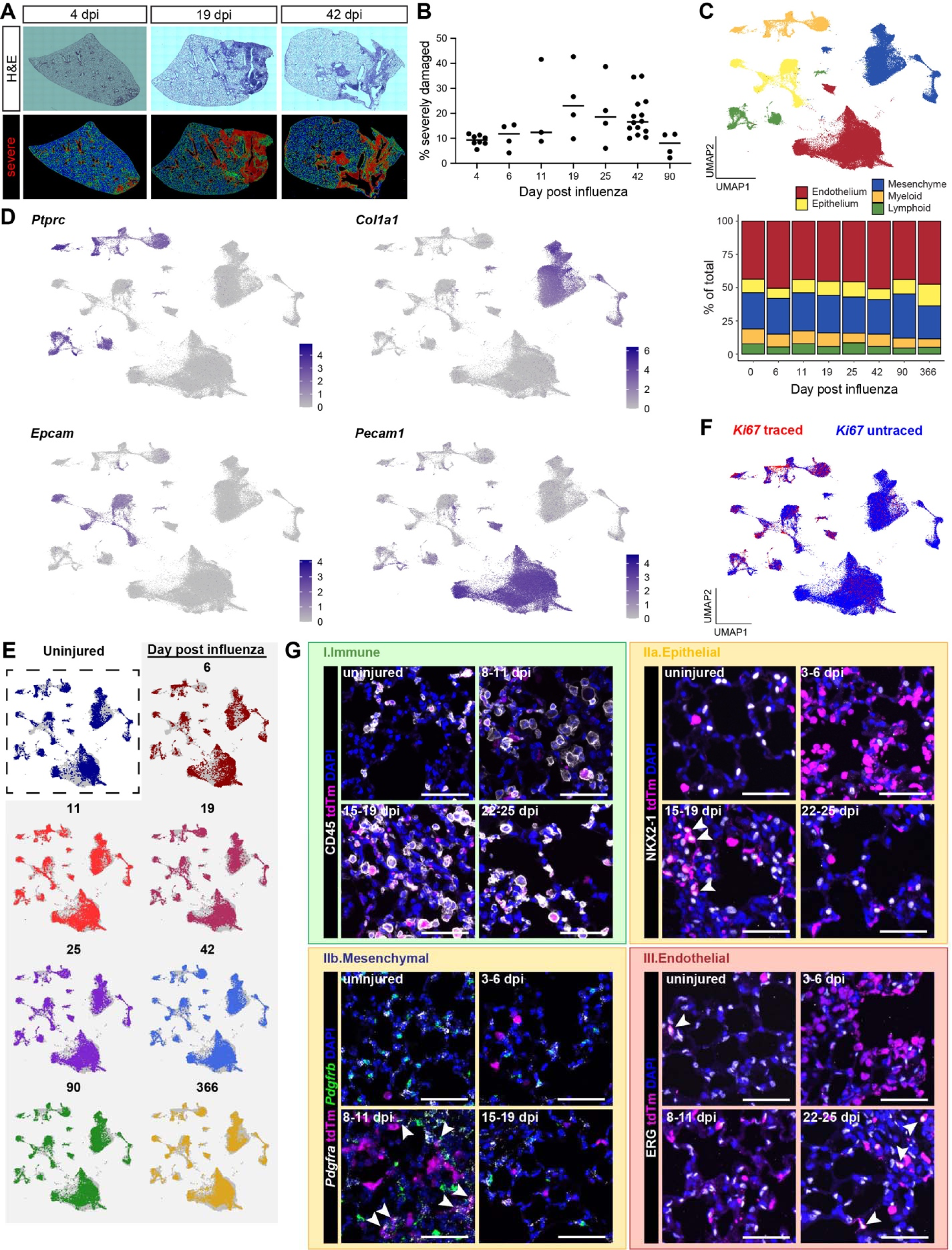
Proliferation occurs in phases across regeneration. **(A)** Representative images of H&E-stained lung tissue sections and output of the computational lung damage assessment program^17^. **(B)** Quantification of the percentage of tissue identified as severely damaged (red) using the lung damage assessment program. **(C)** Merged scRNA-seq data from all time points colored by cellular compartment with proportional composition summarized in bar plot. **(D)** Feature plots for immune (*Ptprc*), mesenchymal (*Col1a1*), epithelial (*Epcam*), and endothelial (*Pecam1*) lineage markers. **(E)** Merged data were evenly down sampled by experimental day and highlighted in the merged UMAP space. **(F)** Traced and untraced cells. **(G)** Representative images from mice used in the scRNA-seq experiment at time points not shown Fig. 2: immune cells (tdTm^+^/CD45^+^) in uninjured mice and 8-11, 15-19, and 22-25 dpi; epithelial cells (tdTm^+^/NKX2-1^+^) in uninjured mice and 3-6, 15-19, and 22-25 dpi; mesenchymal cells (*Pdgfra*^+^ or *Pdgfrb*^+^) in uninjured mice and 3-6, 8-11, and 15-19 dpi; endothelial cells (tdTm^+^/ERG^+^) in uninjured mice and 3-6, 8-11, and 22-25 dpi. Arrow heads demonstrate lineage traced cells for a given cellular compartment. Scale bars 50 μm.

**Figure S3.**
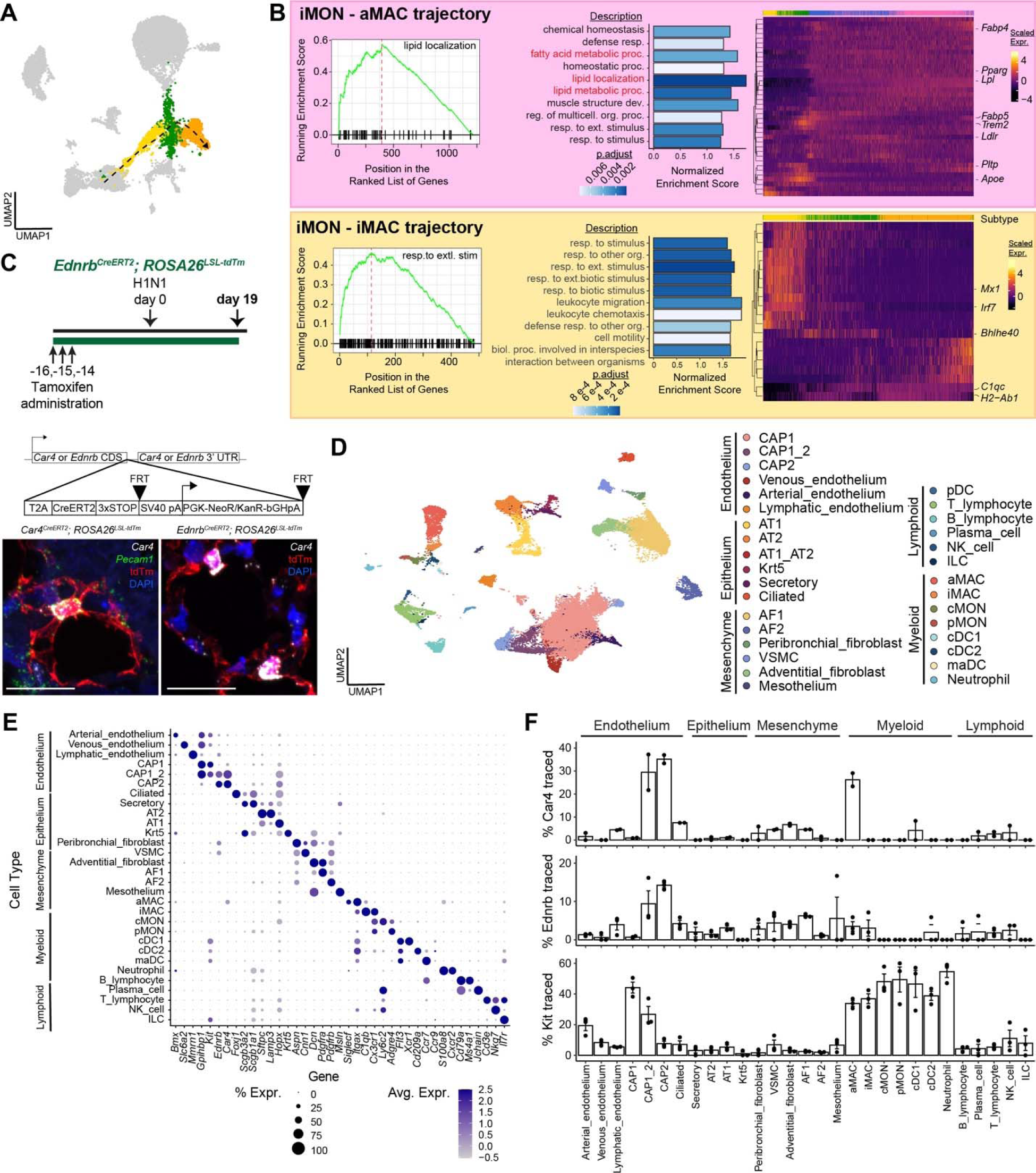
scRNA-seq and lineage tracing of myeloid immune cells or pulmonary endothelial cells using *Kit^MerCreMer^*, *Car4^CreERT2^*, and *Ednrb^CreERT2^ mouse lines.* **(A)** Pseudotime trajectory originating with iMON cells and ending with iMAC cells. **(B)** Gene set enrichment analysis (GSEA) for variable genes along the pseudotime axes of differentiation for aMACs and iMACs. (Left) Running enrichment score for the top hit in each trajectory. (Middle) Normalized enrichment score and p values for the top 10 enriched pathways. (Right) Heatmaps for the top enriched genes in pathways shown at left. **(C)** Experimental schematic for the *Ednrb^CreERT2^* lineage trace experiment and schematic for creation of the *Car4^CreERT2^* and *Ednrb^CreERT2^* mouse lines. Scale bars 25 μm. **(D)** UMAP projection of pooled cells from the Kit, Car4, and Ednrb lineage trace experiment colored by cell type. **(E)** Marker genes expression for cell types shown in (D). **(F)** Percentage of lineage traced cells in each cell type. Each dot represents data from an individual mouse.

**Figure S4.**
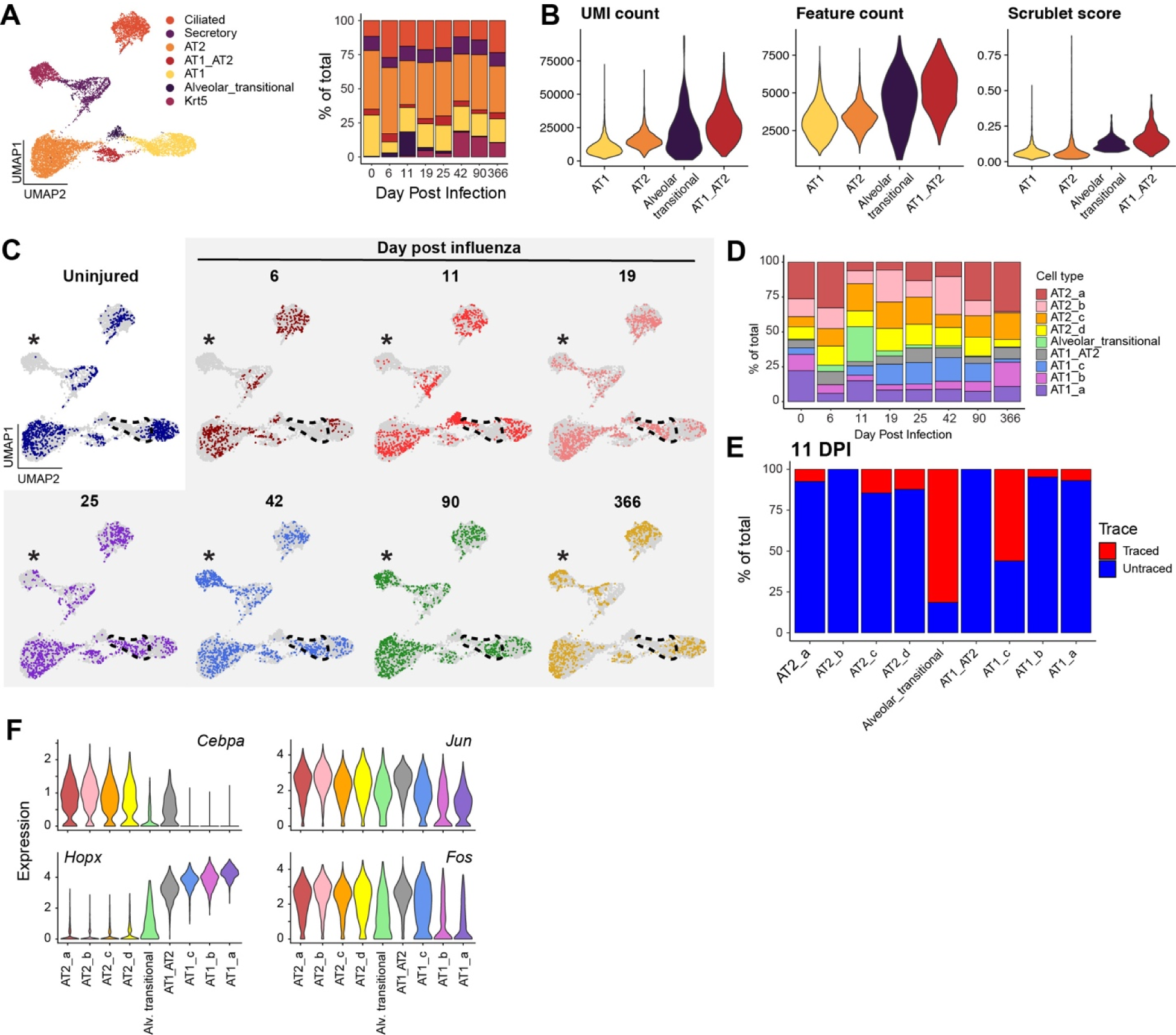
Identification of alveolar cell subtypes and subtype-specific expression patterns. **(A)** All epithelial cells, reclustered at the lineage level and summarized by experimental day. **(B)** UMI count, feature count, and scrublet score for alveolar epithelial cell types. **(C)** Re-clustered cells in the alveolar epithelial compartment were evenly down sampled by experimental day and highlighted accordingly. Dashed lines denote the approximate boundaries of the AT1_c subtype. Asterisks denote the lack of Krt5+ epithelial cells at homeostasis compared to later time points. **(D)** Composition of the alveolar epithelium by experimental day, subset at higher resolution. **(E)** Ki67-tracing of re-clustered alveolar epithelial subtypes at 11 dpi. **(F)** Select transcription factors whose expression is validated in AT2 and AT1 cells (*Cebpa* and *Hopx*), as well as those whose expression decreases between the AT1_c and AT1_a states (*Fos* and *Jun*).

**Figure S5.**
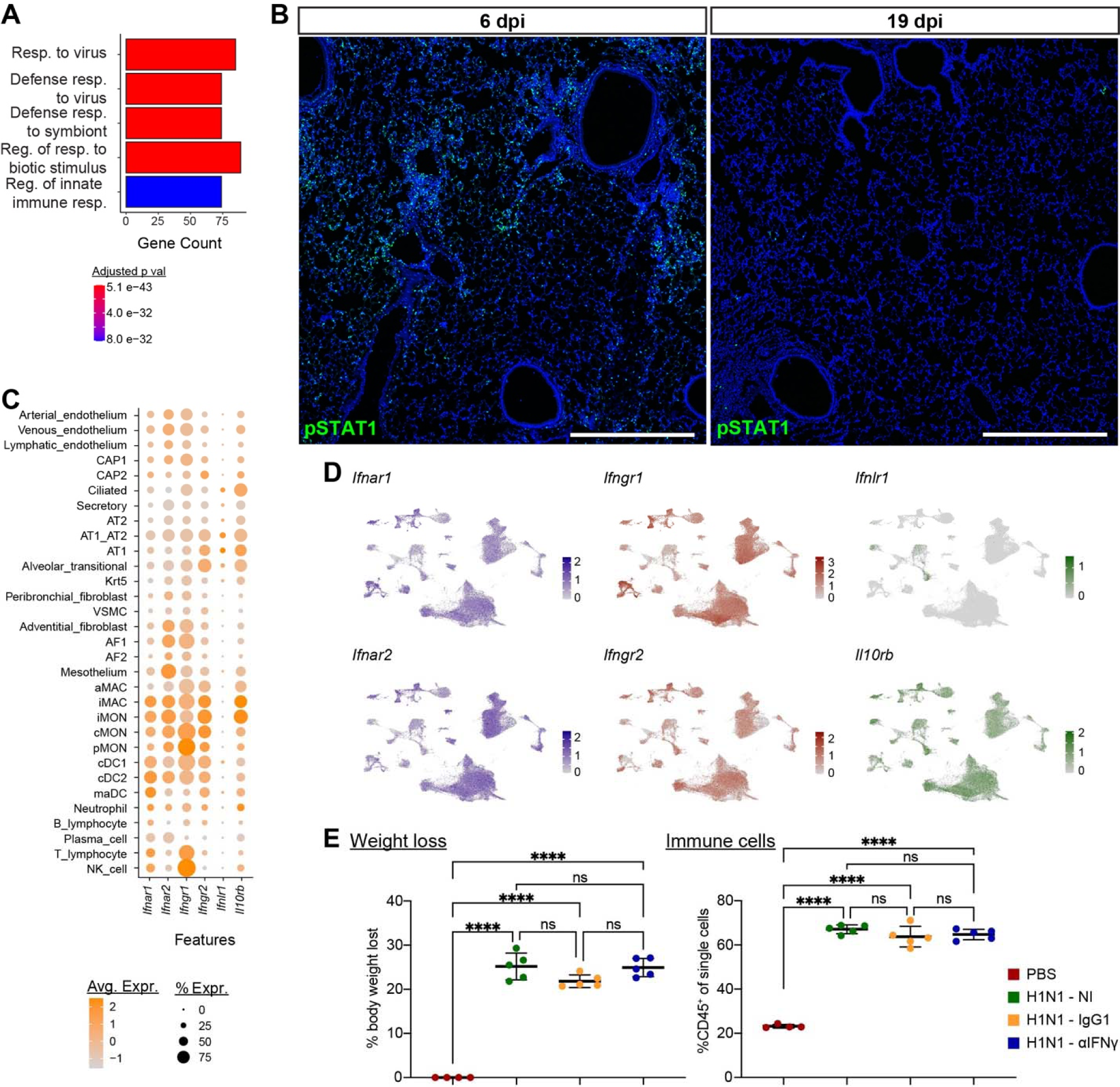
Interferon signaling across regeneration. **(A)** Marker genes were identified for CAP1 cells using experimental day as the grouping variable. The list of differentially expressed genes at 6 dpi was analyzed using GO enrichment. The top 5 enriched categories are shown, colored according to p value. **(B)** Low-magnification image of the distal lung showing pSTAT1 signal, distributed widely throughout the alveolar space at 6 dpi and nearly absent at 19 dpi. Image brightness was adjusted in the low magnification image to improve visualization of fluorescent cells. Scale bars, 500 μm. **(C)** Expression of canonical interferon receptors in all cells of the Ki67 lineage trace dataset. **(D)** UMAP feature plots for data shown in (C). **(E)** Percent body weight lost and flow cytometric quantitation of immune cells in mice from the IFN-γ blocking experiment.

**Figure S6.**
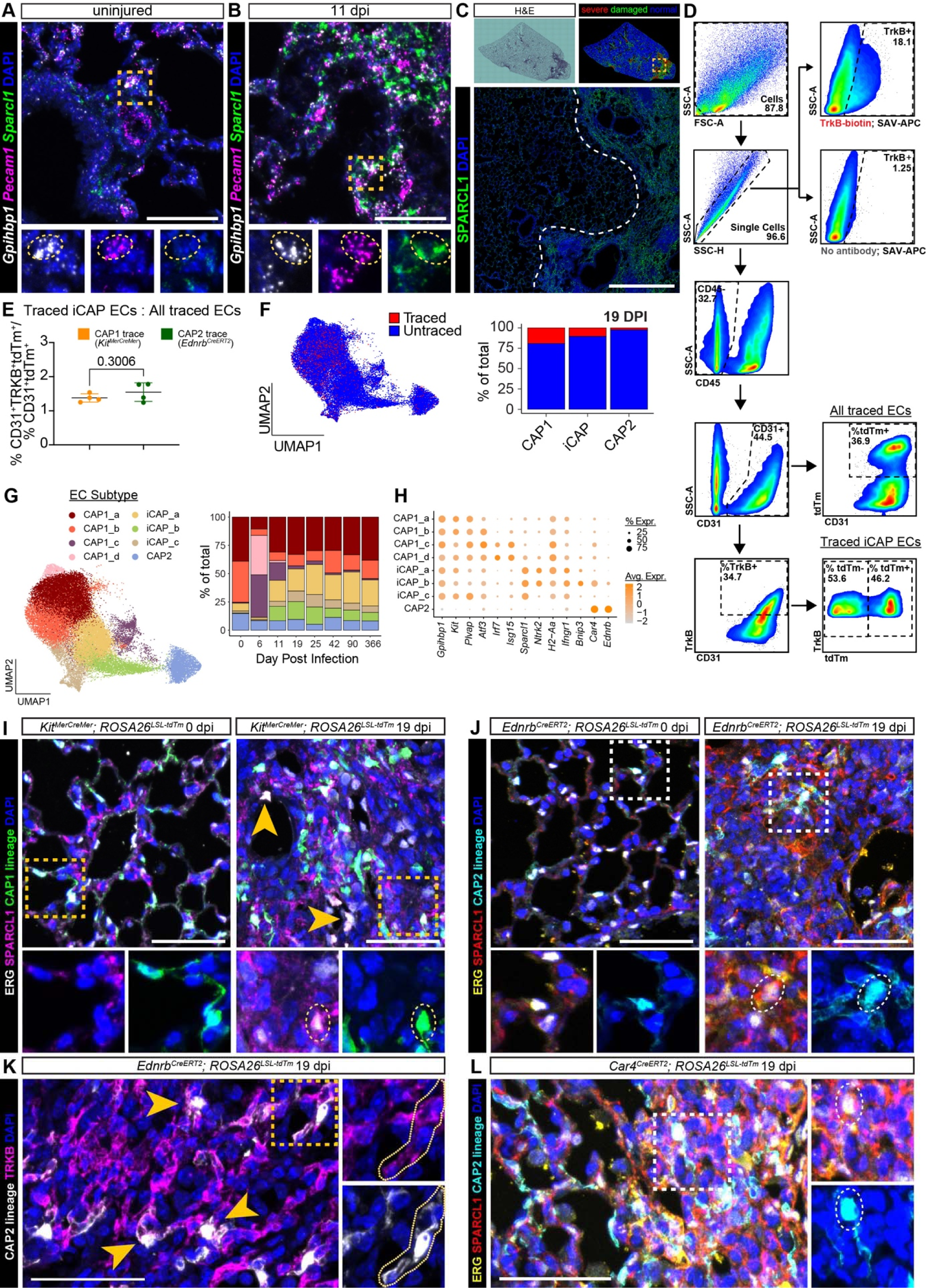
Injury-induced capillary ECs derived from the CAP1 and CAP2 lineages express Sparcl1 and are associated with damaged regions of the lung. RNAscope for *Sparcl1* and CAP1 markers at **(A)** homeostasis and **(B)** 11 dpi. Scale bar 50 μm. **(C)** (Upper) Representative H&E-stained image of a lung lobe at 19 dpi, along with injury severity quantified using a lung damage assessment algorithm^17^. (Lower) Sparcl1 staining in a region spanning injured and uninjured lung. Scale bar 500 μm. **(D)** Flow cytometry gating strategy for identifying traced and untraced TrkB^+^ ECs. Upper right plots demonstrate the specificity of TrkB detection using biotinylated anti-TrkB and APC-conjugated streptavidin (APC-SAV). **(E)** The proportion of iCAP ECs arising from CAP1s or CAP2s was summarized from flow cytometric quantitation and normalized to the fraction of traced ECs. **(F)** Informatically detected Ki67 lineage tracing of capillary ECs types at 19 dpi. **(G)** (Left panel) All EC subtypes identified following re-clustering, including those subtypes not highlighted in Fig 6F. (Right panel) Quantification of each EC subtype, summarized by experimental day. **(H)** Dot plot of select marker genes for each EC subtype. **(I)** IF for Sparcl1 in conjunction with the EC marker ERG and CAP1 lineage trace in *Kit^MerCreMer^*; *ROSA26^LSL-tdTm^*mice at homeostasis and 19 dpi. Scale bars 50 μm. **(J)** IF for Sparcl1 in conjunction with the EC marker ERG and CAP2 lineage trace in *Ednrb^CreERT2^*; *ROSA26^LSL-tdTm^* mice at homeostasis and 19 dpi. Scale bars 50 μm. **(K)** IF for TrkB and the CAP2 lineage trace in *Ednrb^CreERT2^*; *ROSA26^LSL-tdTm^* mice at 19 dpi. Scale bars 50 μm. **(L)** IF for Sparcl1 in conjunction with the EC marker ERG and CAP2 lineage trace in *Car4^CreERT2^*; *ROSA26^LSL-tdTm^* mice at homeostasis and 19 dpi. Scale bars 50 μm.

**Figure S7.**
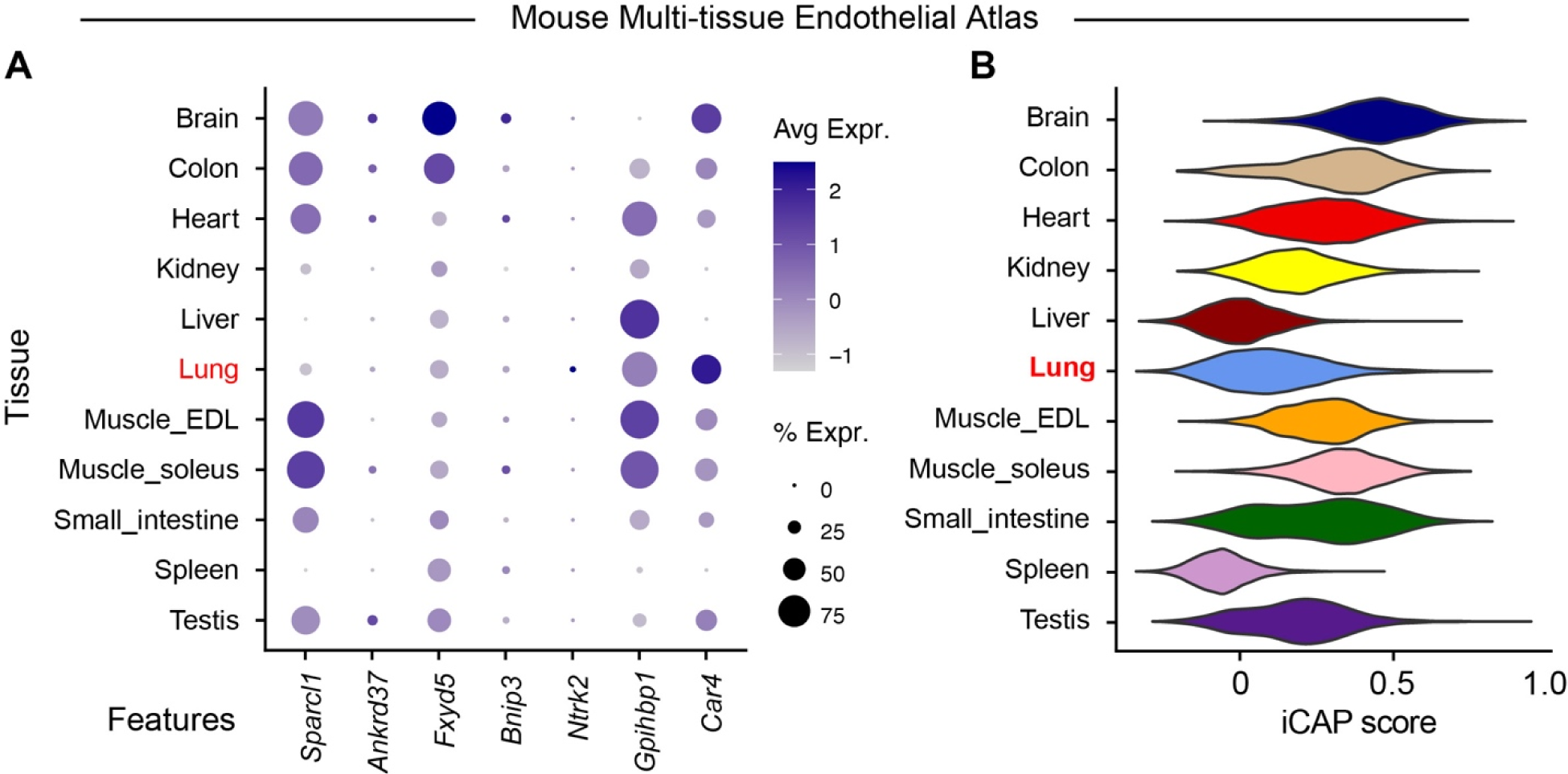
The iCAP signature signifies a non-lung state and is perturbed in non-resolving human viral lung injury. **(A)** Expression of iCAP signature genes in pooled endothelial cells from a mouse whole body endothelial cell atlas (lung highlighted in red). **(B)** A module score incorporating the top 50 iCAP signature genes was generated for all cells in the data set and binned per tissue of origin.

## References

1. Weibel, E.R. (2017). Lung morphometry: the link between structure and function. Cell Tissue Res. 367, 413–426. 10.1007/s00441-016-2541-4.

2. Cao, S., Feng, H., Yi, H., Pan, M., Lin, L., Zhang, Y.S., Feng, Z., Liang, W., Cai, B., Li, Q., et al. (2023). Single-cell RNA sequencing reveals the developmental program underlying proximal-distal patterning of the human lung at the embryonic stage. Cell Res. 33, 421–433. 10.1038/s41422-023-00802-6.

3. He, P., Lim, K., Sun, D., Pett, J.P., Jeng, Q., Polanski, K., Dong, Z., Bolt, L., Richardson, L., Mamanova, L., et al. (2022). A human fetal lung cell atlas uncovers proximal-distal gradients of differentiation and key regulators of epithelial fates. Cell 185, 4841–4860.e25. 10.1016/j.cell.2022.11.005.

4. Miller, A.J., Yu, Q., Czerwinski, M., Tsai, Y.-H., Conway, R.F., Wu, A., Holloway, E.M., Walker, T., Glass, I.A., Treutlein, B., et al. (2020). In Vitro and In Vivo Development of the Human Airway at Single-Cell Resolution. Dev. Cell 53, 117–128.e6. 10.1016/j.devcel.2020.01.033.

5. Negretti, N.M., Plosa, E.J., Benjamin, J.T., Schuler, B.A., Habermann, A.C., Jetter, C.S., Gulleman, P., Bunn, C., Hackett, A.N., Ransom, M., et al. (2021). A single-cell atlas of mouse lung development. Dev. Camb. Engl. 148, dev199512. 10.1242/dev.199512.

6. Zepp, J.A., Morley, M.P., Loebel, C., Kremp, M.M., Chaudhry, F.N., Basil, M.C., Leach, J.P., Liberti, D.C., Niethamer, T.K., Ying, Y., et al. (2021). Genomic, epigenomic, and biophysical cues controlling the emergence of the lung alveolus. Science 371, eabc3172. 10.1126/science.abc3172.

7. Gillich, A., Zhang, F., Farmer, C.G., Travaglini, K.J., Tan, S.Y., Gu, M., Zhou, B., Feinstein, J.A., Krasnow, M.A., and Metzger, R.J. (2020). Capillary cell-type specialization in the alveolus. Nature 586, 785–789. 10.1038/s41586-020-2822-7.

8. Martins, L.R., Glimm, H., and Scholl, C. (2023). Single-cell RNA sequencing of mouse lower respiratory tract epithelial cells: A meta-analysis. Cells Dev. 174, 203847. 10.1016/j.cdev.2023.203847.

9. Niethamer, T.K., Stabler, C.T., Leach, J.P., Zepp, J.A., Morley, M.P., Babu, A., Zhou, S., and Morrisey, E.E. (2020). Defining the role of pulmonary endothelial cell heterogeneity in the response to acute lung injury. eLife 9, e53072. 10.7554/eLife.53072.

10. Paris, A.J., Hayer, K.E., Oved, J.H., Avgousti, D.C., Toulmin, S.A., Zepp, J.A., Zacharias, W.J., Katzen, J.B., Basil, M.C., Kremp, M.M., et al. (2020). STAT3-BDNF-TrkB signalling promotes alveolar epithelial regeneration after lung injury. Nat. Cell Biol. 22, 1197–1210. 10.1038/s41556-020-0569-x.

11. Sikkema, L., Ramírez-Suástegui, C., Strobl, D.C., Gillett, T.E., Zappia, L., Madissoon, E., Markov, N.S., Zaragosi, L.-E., Ji, Y., Ansari, M., et al. (2023). An integrated cell atlas of the lung in health and disease. Nat. Med. 29, 1563–1577. 10.1038/s41591-023-02327-2.

12. Adamson, I.Y., and Bowden, D.H. (1974). The type 2 cell as progenitor of alveolar epithelial regeneration. A cytodynamic study in mice after exposure to oxygen. Lab. Investig. J. Tech. Methods Pathol. 30, 35–42.

13. Barkauskas, C.E., Cronce, M.J., Rackley, C.R., Bowie, E.J., Keene, D.R., Stripp, B.R., Randell, S.H., Noble, P.W., and Hogan, B.L.M. (2013). Type 2 alveolar cells are stem cells in adult lung. J. Clin. Invest. 123, 3025–3036. 10.1172/JCI68782.

14. Basil, M.C., Katzen, J., Engler, A.E., Guo, M., Herriges, M.J., Kathiriya, J.J., Windmueller, R., Ysasi, A.B., Zacharias, W.J., Chapman, H.A., et al. (2020). The Cellular and Physiological Basis for Lung Repair and Regeneration: Past, Present, and Future. Cell Stem Cell 26, 482–502. 10.1016/j.stem.2020.03.009.

15. Hogan, B.L.M., Barkauskas, C.E., Chapman, H.A., Epstein, J.A., Jain, R., Hsia, C.C.W., Niklason, L., Calle, E., Le, A., Randell, S.H., et al. (2014). Repair and regeneration of the respiratory system: complexity, plasticity, and mechanisms of lung stem cell function. Cell Stem Cell 15, 123–138. 10.1016/j.stem.2014.07.012.

16. Cardenas-Diaz, F.L., Liberti, D.C., Leach, J.P., Babu, A., Barasch, J., Shen, T., Diaz-Miranda, M.A., Zhou, S., Ying, Y., Callaway, D.A., et al. (2023). Temporal and spatial staging of lung alveolar regeneration is determined by the grainyhead transcription factor Tfcp2l1. Cell Rep. 42. 10.1016/j.celrep.2023.112451.

17. Liberti, D.C., Kremp, M.M., Liberti, W.A., Penkala, I.J., Li, S., Zhou, S., and Morrisey, E.E. (2021). Alveolar epithelial cell fate is maintained in a spatially restricted manner to promote lung regeneration after acute injury. Cell Rep. 35, 109092. 10.1016/j.celrep.2021.109092.

18. Zacharias, W.J., Frank, D.B., Zepp, J.A., Morley, M.P., Alkhaleel, F.A., Kong, J., Zhou, S., Cantu, E., and Morrisey, E.E. (2018). Regeneration of the lung alveolus by an evolutionarily conserved epithelial progenitor. Nature 555, 251–255. 10.1038/nature25786.

19. Niethamer, T.K., Levin, L.I., Morley, M.P., Babu, A., Zhou, S., and Morrisey, E.E. (2023). Atf3 defines a population of pulmonary endothelial cells essential for lung regeneration. eLife 12, e83835. 10.7554/eLife.83835.

20. Zhao, G., Weiner, A.I., Neupauer, K.M., de Mello Costa, M.F., Palashikar, G., Adams-Tzivelekidis, S., Mangalmurti, N.S., and Vaughan, A.E. (2020). Regeneration of the pulmonary vascular endothelium after viral pneumonia requires COUP-TF2. Sci. Adv. 6, eabc4493. 10.1126/sciadv.abc4493.

21. Choi, J., Park, J.-E., Tsagkogeorga, G., Yanagita, M., Koo, B.-K., Han, N., and Lee, J.-H. (2020). Inflammatory Signals Induce AT2 Cell-Derived Damage-Associated Transient Progenitors that Mediate Alveolar Regeneration. Cell Stem Cell 27, 366–382.e7. 10.1016/j.stem.2020.06.020.

22. Kobayashi, Y., Tata, A., Konkimalla, A., Katsura, H., Lee, R.F., Ou, J., Banovich, N.E., Kropski, J.A., and Tata, P.R. (2020). Persistence of a regeneration-associated, transitional alveolar epithelial cell state in pulmonary fibrosis. Nat. Cell Biol. 22, 934–946. 10.1038/s41556-020-0542-8.

23. Strunz, M., Simon, L.M., Ansari, M., Kathiriya, J.J., Angelidis, I., Mayr, C.H., Tsidiridis, G., Lange, M., Mattner, L.F., Yee, M., et al. (2020). Alveolar regeneration through a Krt8+ transitional stem cell state that persists in human lung fibrosis. Nat. Commun. 11, 3559. 10.1038/s41467-020-17358-3.

24. Katsura, H., Kobayashi, Y., Tata, P.R., and Hogan, B.L.M. (2019). IL-1 and TNFα Contribute to the Inflammatory Niche to Enhance Alveolar Regeneration. Stem Cell Rep. 12, 657–666. 10.1016/j.stemcr.2019.02.013.

25. Lechner, A.J., Driver, I.H., Lee, J., Conroy, C.M., Nagle, A., Locksley, R.M., and Rock, J.R. (2017). Recruited Monocytes and Type 2 Immunity Promote Lung Regeneration following Pneumonectomy. Cell Stem Cell 21, 120–134.e7. 10.1016/j.stem.2017.03.024.

26. Boyd, D.F., Allen, E.K., Randolph, A.G., Guo, X.J., Weng, Y., Sanders, C.J., Bajracharya, R., Lee, N.K., Guy, C.S., Vogel, P., et al. (2020). Exuberant fibroblast activity compromises lung function via ADAMTS4. Nature 587, 466–471. 10.1038/s41586-020-2877-5.

27. Flerlage, T., Boyd, D.F., Meliopoulos, V., Thomas, P.G., and Schultz-Cherry, S. (2021). Influenza virus and SARS-CoV-2: pathogenesis and host responses in the respiratory tract. Nat. Rev. Microbiol. 19, 425–441. 10.1038/s41579-021-00542-7.

28. Major, J., Crotta, S., Llorian, M., McCabe, T.M., Gad, H.H., Priestnall, S.L., Hartmann, R., and Wack, A. (2020). Type I and III interferons disrupt lung epithelial repair during recovery from viral infection. Science 369, 712–717. 10.1126/science.abc2061.

29. Basak, O., Krieger, T.G., Muraro, M.J., Wiebrands, K., Stange, D.E., Frias-Aldeguer, J., Rivron, N.C., van de Wetering, M., van Es, J.H., van Oudenaarden, A., et al. (2018). Troy+ brain stem cells cycle through quiescence and regulate their number by sensing niche occupancy. Proc. Natl. Acad. Sci. U. S. A. 115, E610–E619. 10.1073/pnas.1715911114.

30. Madisen, L., Zwingman, T.A., Sunkin, S.M., Oh, S.W., Zariwala, H.A., Gu, H., Ng, L.L., Palmiter, R.D., Hawrylycz, M.J., Jones, A.R., et al. (2010). A robust and high-throughput Cre reporting and characterization system for the whole mouse brain. Nat. Neurosci. 13, 133–140. 10.1038/nn.2467.

31. Sun, X., Perl, A.-K., Li, R., Bell, S.M., Sajti, E., Kalinichenko, V.V., Kalin, T.V., Misra, R.S., Deshmukh, H., Clair, G., et al. (2022). A census of the lung: CellCards from LungMAP. Dev. Cell 57, 112–145.e2. 10.1016/j.devcel.2021.11.007.

32. Malainou, C., Peteranderl, C., Matt, U., Vazquez-Armendariz, A.I., Better, J., Schultheis, H., Hoppe, J., Firsching, T., Gruber, A., Guenther, S., et al. (2023). Tnfsf14-driven apoptosis of alveolar macrophages upon influenza infection enables the establishment of secondary pneumococcal pneumonia. ERJ Open Res. 9. 10.1183/23120541.LSC-2023.170.

33. Li, F., Piattini, F., Pohlmeier, L., Feng, Q., Rehrauer, H., and Kopf, M. (2022). Monocyte-derived alveolar macrophages autonomously determine severe outcome of respiratory viral infection. Sci. Immunol. 7, eabj5761. 10.1126/sciimmunol.abj5761.

34. Aegerter, H., Kulikauskaite, J., Crotta, S., Patel, H., Kelly, G., Hessel, E.M., Mack, M., Beinke, S., and Wack, A. (2020). Influenza-induced monocyte-derived alveolar macrophages confer prolonged antibacterial protection. Nat. Immunol. 21, 145–157. 10.1038/s41590-019-0568-x.

35. Li, F., Okreglicka, K.M., Pohlmeier, L.M., Schneider, C., and Kopf, M. (2020). Fetal monocytes possess increased metabolic capacity and replace primitive macrophages in tissue macrophage development. EMBO J. 39, e103205. 10.15252/embj.2019103205.

36. Misharin, A.V., Morales-Nebreda, L., Reyfman, P.A., Cuda, C.M., Walter, J.M., McQuattie-Pimentel, A.C., Chen, C.-I., Anekalla, K.R., Joshi, N., Williams, K.J.N., et al. (2017). Monocyte-derived alveolar macrophages drive lung fibrosis and persist in the lung over the life span. J. Exp. Med. 214, 2387–2404. 10.1084/jem.20162152.

37. Yao, Y., Jeyanathan, M., Haddadi, S., Barra, N.G., Vaseghi-Shanjani, M., Damjanovic, D., Lai, R., Afkhami, S., Chen, Y., Dvorkin-Gheva, A., et al. (2018). Induction of Autonomous Memory Alveolar Macrophages Requires T Cell Help and Is Critical to Trained Immunity. Cell 175, 1634–1650.e17. 10.1016/j.cell.2018.09.042.

38. Aegerter, H., Lambrecht, B.N., and Jakubzick, C.V. (2022). Biology of lung macrophages in health and disease. Immunity 55, 1564–1580. 10.1016/j.immuni.2022.08.010.

39. Fabre, T., Barron, A.M.S., Christensen, S.M., Asano, S., Bound, K., Lech, M.P., Wadsworth, M.H., Chen, X., Wang, C., Wang, J., et al. (2023). Identification of a broadly fibrogenic macrophage subset induced by type 3 inflammation. Sci. Immunol. 8, eadd8945. 10.1126/sciimmunol.add8945.

40. Hoeft, K., Schaefer, G.J.L., Kim, H., Schumacher, D., Bleckwehl, T., Long, Q., Klinkhammer, B.M., Peisker, F., Koch, L., Nagai, J., et al. (2023). Platelet-instructed SPP1+ macrophages drive myofibroblast activation in fibrosis in a CXCL4-dependent manner. Cell Rep. 42, 112131. 10.1016/j.celrep.2023.112131.

41. Takahashi, F., Takahashi, K., Shimizu, K., Cui, R., Tada, N., Takahashi, H., Soma, S., Yoshioka, M., and Fukuchi, Y. (2004). Osteopontin is strongly expressed by alveolar macrophages in the lungs of acute respiratory distress syndrome. Lung 182, 173–185. 10.1007/s00408-004-0309-1.

42. Maus, U.A., Janzen, S., Wall, G., Srivastava, M., Blackwell, T.S., Christman, J.W., Seeger, W., Welte, T., and Lohmeyer, J. (2006). Resident alveolar macrophages are replaced by recruited monocytes in response to endotoxin-induced lung inflammation. Am. J. Respir. Cell Mol. Biol. 35, 227–235. 10.1165/rcmb.2005-0241OC.

43. Cheong, C., Matos, I., Choi, J.-H., Dandamudi, D.B., Shrestha, E., Longhi, M.P., Jeffrey, K.L., Anthony, R.M., Kluger, C., Nchinda, G., et al. (2010). Microbial Stimulation Fully Differentiates Monocytes to DC-SIGN/CD209+ Dendritic Cells for Immune T Cell Areas. Cell 143, 416–429. 10.1016/j.cell.2010.09.039.

44. Serbina, N.V., Salazar-Mather, T.P., Biron, C.A., Kuziel, W.A., and Pamer, E.G. (2003). TNF/iNOS-Producing Dendritic Cells Mediate Innate Immune Defense against Bacterial Infection. Immunity 19, 59–70. 10.1016/S1074-7613(03)00171-7.

45. Sabatel, C., Radermecker, C., Fievez, L., Paulissen, G., Chakarov, S., Fernandes, C., Olivier, S., Toussaint, M., Pirottin, D., Xiao, X., et al. (2017). Exposure to Bacterial CpG DNA Protects from Airway Allergic Inflammation by Expanding Regulatory Lung Interstitial Macrophages. Immunity 46, 457–473. 10.1016/j.immuni.2017.02.016.

46. Tan, S.Y.S., and Krasnow, M.A. (2016). Developmental origin of lung macrophage diversity. Dev. Camb. Engl. 143, 1318–1327. 10.1242/dev.129122.

47. Schneider, C., Nobs, S.P., Kurrer, M., Rehrauer, H., Thiele, C., and Kopf, M. (2014). Induction of the nuclear receptor PPAR-γ by the cytokine GM-CSF is critical for the differentiation of fetal monocytes into alveolar macrophages. Nat. Immunol. 15, 1026–1037. 10.1038/ni.3005.

48. Sajti, E., Link, V.M., Ouyang, Z., Spann, N.J., Westin, E., Romanoski, C.E., Fonseca, G.J., Prince, L.S., and Glass, C.K. (2020). Transcriptomic and epigenetic mechanisms underlying myeloid diversity in the lung. Nat. Immunol. 21, 221–231. 10.1038/s41590-019-0582-z.

49. van Berlo, J.H., Kanisicak, O., Maillet, M., Vagnozzi, R.J., Karch, J., Lin, S.-C.J., Middleton, R.C., Marbán, E., and Molkentin, J.D. (2014). c-kit+ cells minimally contribute cardiomyocytes to the heart. Nature 509, 337–341. 10.1038/nature13309.

50. Shen, W.-K., Chen, S.-Y., Gan, Z.-Q., Zhang, Y.-Z., Yue, T., Chen, M.-M., Xue, Y., Hu, H., and Guo, A.-Y. (2023). AnimalTFDB 4.0: a comprehensive animal transcription factor database updated with variation and expression annotations. Nucleic Acids Res. 51, D39–D45. 10.1093/nar/gkac907.

51. Daniel, B., Nagy, G., Czimmerer, Z., Horvath, A., Hammers, D.W., Cuaranta-Monroy, I., Poliska, S., Tzerpos, P., Kolostyak, Z., Hays, T.T., et al. (2018). The Nuclear Receptor PPARγ Controls Progressive Macrophage Polarization as a Ligand-Insensitive Epigenomic Ratchet of Transcriptional Memory. Immunity 49, 615–626.e6. 10.1016/j.immuni.2018.09.005.

52. Rauschmeier, R., Gustafsson, C., Reinhardt, A., A-Gonzalez, N., Tortola, L., Cansever, D., Subramanian, S., Taneja, R., Rossner, M.J., Sieweke, M.H., et al. (2019). Bhlhe40 and Bhlhe41 transcription factors regulate alveolar macrophage self-renewal and identity. EMBO J. 38, e101233. 10.15252/embj.2018101233.

53. Vanneste, D., Bai, Q., Hasan, S., Peng, W., Pirottin, D., Schyns, J., Maréchal, P., Ruscitti, C., Meunier, M., Liu, Z., et al. (2023). MafB-restricted local monocyte proliferation precedes lung interstitial macrophage differentiation. Nat. Immunol. 24, 827–840. 10.1038/s41590-023-01468-3.

54. Jin, S., Guerrero-Juarez, C.F., Zhang, L., Chang, I., Ramos, R., Kuan, C.-H., Myung, P., Plikus, M.V., and Nie, Q. (2021). Inference and analysis of cell-cell communication using CellChat. Nat. Commun. 12, 1088. 10.1038/s41467-021-21246-9.

55. Quinlan, G.J., Lamb, N.J., Evans, T.W., and Gutteridge, J.M. (1996). Plasma fatty acid changes and increased lipid peroxidation in patients with adult respiratory distress syndrome. Crit. Care Med. 24, 241–246. 10.1097/00003246-199602000-00010.

56. Al-Saiedy, M., Gunasekara, L., Green, F., Pratt, R., Chiu, A., Yang, A., Dennis, J., Pieron, C., Bjornson, C., Winston, B., et al. (2018). Surfactant Dysfunction in ARDS and Bronchiolitis is Repaired with Cyclodextrins. Mil. Med. 183, 207–215. 10.1093/milmed/usx204.

57. Toth, A., Kannan, P., Snowball, J., Kofron, M., Wayman, J.A., Bridges, J.P., Miraldi, E.R., Swarr, D., and Zacharias, W.J. (2023). Alveolar epithelial progenitor cells require Nkx2-1 to maintain progenitor-specific epigenomic state during lung homeostasis and regeneration. Nat. Commun. 14, 8452. 10.1038/s41467-023-44184-0.

58. Basil, M.C., Cardenas-Diaz, F.L., Kathiriya, J.J., Morley, M.P., Carl, J., Brumwell, A.N., Katzen, J., Slovik, K.J., Babu, A., Zhou, S., et al. (2022). Human distal airways contain a multipotent secretory cell that can regenerate alveoli. Nature 604, 120–126. 10.1038/s41586-022-04552-0.

59. Fernanda de Mello Costa, M., Weiner, A.I., and Vaughan, A.E. (2020). Basal-like Progenitor Cells: A Review of Dysplastic Alveolar Regeneration and Remodeling in Lung Repair. Stem Cell Rep. 15, 1015–1025. 10.1016/j.stemcr.2020.09.006.

60. Vaughan, A.E., Brumwell, A.N., Xi, Y., Gotts, J.E., Brownfield, D.G., Treutlein, B., Tan, K., Tan, V., Liu, F.C., Looney, M.R., et al. (2015). Lineage-negative progenitors mobilize to regenerate lung epithelium after major injury. Nature 517, 621–625. 10.1038/nature14112.

61. Joseph, R., Dou, D., and Tsang, W. (1995). Neuronatin mRNA: alternatively spliced forms of a novel brain-specific mammalian developmental gene. Brain Res. 690, 92–98. 10.1016/0006-8993(95)00621-V.

62. Cimino, I., Rimmington, D., Tung, Y.C.L., Lawler, K., Larraufie, P., Kay, R.G., Virtue, S., Lam, B.Y.H., Fagnocchi, L., Ma, M.K.L., et al. (2021). Murine neuronatin deficiency is associated with a hypervariable food intake and bimodal obesity. Sci. Rep. 11, 17571. 10.1038/s41598-021-96278-8.

63. Wang, Y., Tang, Z., Huang, H., Li, J., Wang, Z., Yu, Y., Zhang, C., Li, J., Dai, H., Wang, F., et al. (2018). Pulmonary alveolar type I cell population consists of two distinct subtypes that differ in cell fate. Proc. Natl. Acad. Sci. U. S. A. 115, 2407–2412. 10.1073/pnas.1719474115.

64. Vila Ellis, L., Cain, M.P., Hutchison, V., Flodby, P., Crandall, E.D., Borok, Z., Zhou, B., Ostrin, E.J., Wythe, J.D., and Chen, J. (2020). Epithelial Vegfa Specifies a Distinct Endothelial Population in the Mouse Lung. Dev. Cell 52, 617–630.e6. 10.1016/j.devcel.2020.01.009.

65. Toulmin, S.A., Bhadiadra, C., Paris, A.J., Lin, J.H., Katzen, J., Basil, M.C., Morrisey, E.E., Worthen, G.S., and Eisenlohr, L.C. (2021). Type II alveolar cell MHCII improves respiratory viral disease outcomes while exhibiting limited antigen presentation. Nat. Commun. 12. 10.1038/s41467-021-23619-6.

66. Zhang, Z., Newton, K., Kummerfeld, S.K., Webster, J., Kirkpatrick, D.S., Phu, L., Eastham-Anderson, J., Liu, J., Lee, W.P., Wu, J., et al. (2017). Transcription factor Etv5 is essential for the maintenance of alveolar type II cells. Proc. Natl. Acad. Sci. 114, 3903–3908. 10.1073/pnas.1621177114.

67. Little, D.R., Lynch, A.M., Yan, Y., Akiyama, H., Kimura, S., and Chen, J. (2021). Differential chromatin binding of the lung lineage transcription factor NKX2-1 resolves opposing murine alveolar cell fates in vivo. Nat. Commun. 12, 2509. 10.1038/s41467-021-22817-6.

68. Gokey, J.J., Snowball, J., Sridharan, A., Sudha, P., Kitzmiller, J.A., Xu, Y., and Whitsett, J.A. (2021). YAP regulates alveolar epithelial cell differentiation and AGER via NFIB/KLF5/NKX2-1. iScience 24, 102967. 10.1016/j.isci.2021.102967.

69. Mostafavi, S., Yoshida, H., Moodley, D., LeBoité, H., Rothamel, K., Raj, T., Ye, C.J., Chevrier, N., Zhang, S.-Y., Feng, T., et al. (2016). Parsing the Interferon Transcriptional Network and Its Disease Associations. Cell 164, 564–578. 10.1016/j.cell.2015.12.032.

70. Iwasaki, A., and Pillai, P.S. (2014). Innate immunity to influenza virus infection. Nat. Rev. Immunol. 14, 315–328. 10.1038/nri3665.

71. Sadler, A.J., and Williams, B.R.G. (2008). Interferon-inducible antiviral effectors. Nat. Rev. Immunol. 8, 559–568. 10.1038/nri2314.

72. Friesel, R., Komoriya, A., and Maciag, T. (1987). Inhibition of endothelial cell proliferation by gamma-interferon. J. Cell Biol. 104, 689–696. 10.1083/jcb.104.3.689.

73. Baumgarth, N., and Kelso, A. (1996). In vivo blockade of gamma interferon affects the influenza virus-induced humoral and the local cellular immune response in lung tissue. J. Virol. 70, 4411–4418.

74. Zhang, L., Gao, S., White, Z., Dai, Y., Malik, A.B., and Rehman, J. (2022). Single-cell transcriptomic profiling of lung endothelial cells identifies dynamic inflammatory and regenerative subpopulations. JCI Insight 7, e158079. 10.1172/jci.insight.158079.

75. Zhao, G., Gentile, M.E., Xue, L., Cosgriff, C.V., Weiner, A.I., Adams-Tzivelekidis, S., Wong, J., Li, X., Kass-Gergi, S., Holcomb, N.P., et al. (2023). Vascular Endothelial-derived SPARCL1 Exacerbates Viral Pneumonia Through Pro-Inflammatory Macrophage Activation. BioRxiv Prepr. Serv. Biol., 2023.05.25.541966. 10.1101/2023.05.25.541966.

76. Kumar, P.A., Hu, Y., Yamamoto, Y., Hoe, N.B., Wei, T.S., Mu, D., Sun, Y., Joo, L.S., Dagher, R., Zielonka, E.M., et al. (2011). Distal airway stem cells yield alveoli in vitro and during lung regeneration following H1N1 influenza infection. Cell 147, 525–538. 10.1016/j.cell.2011.10.001.

77. Zuo, W., Zhang, T., Wu, D.Z., Guan, S.P., Liew, A.-A., Yamamoto, Y., Wang, X., Lim, S.J., Vincent, M., Lessard, M., et al. (2015). p63(+)Krt5(+) distal airway stem cells are essential for lung regeneration. Nature 517, 616–620. 10.1038/nature13903.

78. Kalucka, J., de Rooij, L.P.M.H., Goveia, J., Rohlenova, K., Dumas, S.J., Meta, E., Conchinha, N.V., Taverna, F., Teuwen, L.-A., Veys, K., et al. (2020). Single-Cell Transcriptome Atlas of Murine Endothelial Cells. Cell 180, 764–779.e20. 10.1016/j.cell.2020.01.015.

79. Raslan, A.A., Pham, T.X., Lee, J., Hong, J., Schmottlach, J., Nicolas, K., Dinc, T., Bujor, A.M., Caporarello, N., Thiriot, A., et al. (2023). Single Cell Transcriptomics of Fibrotic Lungs Unveils Aging-associated Alterations in Endothelial and Epithelial Cell Regeneration. BioRxiv Prepr. Serv. Biol., 2023.01.17.523179. 10.1101/2023.01.17.523179.

80. Polverino, F., Celli, B.R., and Owen, C.A. (2018). COPD as an endothelial disorder: endothelial injury linking lesions in the lungs and other organs? (2017 Grover Conference Series). Pulm. Circ. 8, 2045894018758528. 10.1177/2045894018758528.

81. Clarenbach, C.F., Senn, O., Sievi, N.A., Camen, G., van Gestel, A.J.R., Rossi, V.A., Puhan, M.A., Thurnheer, R., Russi, E.W., and Kohler, M. (2013). Determinants of endothelial function in patients with COPD. Eur. Respir. J. 42, 1194–1204. 10.1183/09031936.00144612.

82. Theodorakopoulou, M.P., Alexandrou, M.E., Bakaloudi, D.R., Pitsiou, G., Stanopoulos, I., Kontakiotis, T., and Boutou, A.K. (2021). Endothelial dysfunction in COPD: a systematic review and meta-analysis of studies using different functional assessment methods. ERJ Open Res. 7, 00983–02020. 10.1183/23120541.00983-2020.

83. Diamond, J.M., Lee, J.C., Kawut, S.M., Shah, R.J., Localio, A.R., Bellamy, S.L., Lederer, D.J., Cantu, E., Kohl, B.A., Lama, V.N., et al. (2013). Clinical risk factors for primary graft dysfunction after lung transplantation. Am. J. Respir. Crit. Care Med. 187, 527–534. 10.1164/rccm.201210-1865OC.

84. Sörensen, I., Adams, R.H., and Gossler, A. (2009). DLL1-mediated Notch activation regulates endothelial identity in mouse fetal arteries. Blood 113, 5680–5688. 10.1182/blood-2008-08-174508.

85. Windmueller, R., Leach, J.P., Babu, A., Zhou, S., Morley, M.P., Wakabayashi, A., Petrenko, N.B., Viatour, P., and Morrisey, E.E. (2020). Direct Comparison of Mononucleated and Binucleated Cardiomyocytes Reveals Molecular Mechanisms Underlying Distinct Proliferative Competencies. Cell Rep. 30, 3105–3116.e4. 10.1016/j.celrep.2020.02.034.

86. Young, M.D., and Behjati, S. (2020). SoupX removes ambient RNA contamination from droplet-based single-cell RNA sequencing data. GigaScience 9, giaa151. 10.1093/gigascience/giaa151.

87. Y, H., S, H., E, A.-N., Wm, M., S, Z., A, B., Mj, L., Aj, W., C, D., M, Z., et al. (2021). Integrated analysis of multimodal single-cell data. Cell 184. 10.1016/j.cell.2021.04.048.

88. Street, K., Risso, D., Fletcher, R.B., Das, D., Ngai, J., Yosef, N., Purdom, E., and Dudoit, S. (2018). Slingshot: cell lineage and pseudotime inference for single-cell transcriptomics. BMC Genomics 19, 477. 10.1186/s12864-018-4772-0.

89. Van den Berge, K., Roux de Bézieux, H., Street, K., Saelens, W., Cannoodt, R., Saeys, Y., Dudoit, S., and Clement, L. (2020). Trajectory-based differential expression analysis for single-cell sequencing data. Nat. Commun. 11, 1–13. 10.1038/s41467-020-14766-3.

